# Ccdc66 regulates primary cilium stability, disassembly and signaling important for epithelial organization

**DOI:** 10.1101/2024.06.16.599243

**Authors:** Jovana Deretic, Seyma Cengiz-Emek, Ece Seyrek, Elif Nur Firat-Karalar

## Abstract

The primary cilium is a conserved, microtubule-based organelle that transduces signaling pathways essential for development and homeostasis. It is a dynamic structure that assembles and disassembles in response to intrinsic and extrinsic stimuli while maintaining remarkable stability and tightly controlled length. Although cilium assembly is well-understood, less is known about the molecular players and pathways governing their stability, length and disassembly. Here, we elucidated the function of Ccdc66, a microtubule-associated protein linked to ciliopathies, in cilium maintenance and disassembly in mouse epithelial cells. We found that Ccdc66 depletion disrupts cilium disassembly, length and stability, but does not affect assembly in these cells. Live imaging of these processes revealed that cilia in Ccdc66-depleted cells frequently fluctuate in length and exhibit increased ectocytosis from the cilium tip. Phenotypic rescue experiments and *in vitro* microtubule stabilization assays showed that Ccdc66 mediates these functions via regulating the stability of microtubules. Temporal proximity mapping of CCDC66 identified potential new regulators and molecular pathways involved in cilium disassembly. Additionally, depletion of CCDC66 compromised Hedgehog and Wnt pathway activation and disrupted epithelial cell organization and polarity in two-dimensional and three-dimensional cultures. Collectively, our results define Ccdc66 as a new microtubule-stabilizing factor that regulates cilium stability and disassembly, providing insights into the mechanisms of cilium homeostasis and the pathologies associated with Ccdc66.

## Introduction

The primary cilium is an evolutionary conserved, microtubule-based cellular projection that transduces signaling pathways essential for development and homeostasis, including Hedgehog and Wnt signaling [1, 2]. It is composed of a microtubule-based axoneme surrounded by a ciliary membrane. The nine radially arranged microtubule doublets of the axoneme are exceptionally stable, mechanically resistant and highly modified by posttranslational modifications (PTMs) [3–6]. The axoneme supports the shape and structure of the cilium and acts as a track for ciliary transport complexes [7, 8]. The ciliary membrane is continuous with the plasma membrane but distinct in its lipid and protein composition. Sensory functions of the cilium require tight spatiotemporal control of its assembly kinetics, length, stability, structure and composition. Deregulation of these controls causes various human diseases including cancer and the multisystem pathologies of the eye, kidney, skeleton, brain and other organs, collectively named “ciliopathies” [9, 10]. Defining the mechanisms by which a functional cilium is assembled, maintained and disassembled in response to different signals and across different cell types is essential to uncover the molecular defects underpinning these diseases.

Primary cilium assembly, maintenance and disassembly are highly regulated processes governed by intrinsic and extrinsic stimuli such as cell cycle cues [8, 11]. In most cells, primary cilium assembles as cells exit the cell cycle during differentiation and quiescence. The assembly process involves the maturation of the mother centriole into a basal body, the elongation of the axoneme from the basal body by Intraflagellar Transport (IFT)-mediated transport of ciliary components and the formation of the ciliary membrane and the ciliary gate [11]. Primary cilium forms via extracellular and intracellular pathways, classified based on whether cilium growth is initiated at the cell surface or within the cytoplasm, respectively [12, 13]. The extracellular pathway is commonly observed in kidney and lung epithelial cells, whereas the intracellular pathway is typical in fibroblasts and retinal epithelial cells.

Once a cilium is formed, it maintains its length, integrity and structure despite the continuous turnover of its lipid and protein composition [4, 14]. Cilium shortening, over-elongation and instability have been reported in ciliopathies, highlighting the physiological significance of maintaining proper cilium length and stability for its functions [10, 15–18]. To date, various mechanisms regulating cilium length and stability have been uncovered through functional screens and targeted molecular and cellular studies. These studies have identified proteins involved in diverse cellular components and functions, including cell cycle regulation, organization of the actin and microtubule cytoskeleton, lipid metabolism, vesicle trafficking (highlighting specific lipids such as PIP2 and processes like endocytosis and ectocytosis), and transcription [19–29]. Within this context, microtubule-associated proteins (MAPs), which regulate microtubule nucleation, dynamics, and stability, have been recognized as critical regulators of these ciliary properties by acting on axonemal microtubules [7, 30]. The best-characterized MAPs for axoneme length control are the molecular motors that cooperate with IFT machinery during transport of ciliary proteins such as tubulin dimers [31]. However, the roles of non-IFT ciliary MAPs beyond molecular motors remain less understood. Therefore, further identification and characterization of these MAPs are necessary to elucidate the mechanisms by which axonemal microtubules are elongated from the basal body to the correct length and how their length and stability are maintained.

Upon mitotic entry, the primary cilium of a quiescent cell disassembles through the resorption of the axoneme, in which the axoneme is depolymerized and ciliary contents are incorporated into the cell. Cilium resorption occurs in two distinct phases [32]. In the first phase, HEF1-Aurora A kinase (AURKA)-histone deacetylase 6 (HDAC6) pathway plays a crucial role [33]. The activation of AURKA by HEF1 leads to the phosphorylation of HDAC6, which then deacetylates axonemal microtubules and destabilizes the axoneme. In addition, the phosphorylation-stimulated activity of KIF2A leads to depolymerization of axonemal microtubules [34]. The second phase involves NEK2-KIF24-dependent microtubule depolymerization at the ciliary base [35]. Live imaging studies of cilium disassembly in mouse kidney epithelial cells have identified an additional disassembly mechanism termed deciliation, where the axoneme is excised near its base [8]. This process is partly facilitated by the microtubule-severing enzyme katanin [36, 37]. Additionally, recent studies in adult differentiating cerebellar granule cells have uncovered a post-mitotic cilium deconstruction pathway involving previously undescribed disassembly intermediates [38, 39]. This discovery reveals a new pathway of cilium disassembly, highlighting its complexity. Despite our comprehensive understanding of the molecular pathways and players involved in primary cilium assembly, little is known about the mechanisms underlying primary cilium disassembly, particularly regarding the role of MAPs in this process.

We previously identified Ccdc66 as a MAP that plays important roles in primary cilium assembly, ciliary signaling and the progression of mitosis and cytokinesis [40–42]. CCDC66 exhibits a dynamic localization profile, ranging from localization to the ciliary axoneme, centrosome and centriolar satellites in quiescent cells, to the mitotic spindle, the central spindle and the midbody in dividing cells [40, 42, 43]. *In vitro* studies showed that CCDC66 directly binds to microtubules and promotes their bundling [40, 41]. Additionally, genetic studies in dogs and mice have linked early frameshift mutations and deletions of CCDC66 to retinal degeneration and olfactory deficits, underscoring its physiological significance [44–47]. Moreover, CCDC66 is part of a ciliary tip module that includes other MAPs such as CEP104, CSPP1, TOGARAM1 and ARMC9, which are mutated in the neurodevelopmental ciliopathy known as Joubert syndrome [48, 49]. A recent *in vitro* reconstitution study showed that these proteins collectively stabilize microtubule plus ends and promote their slow and persistent elongation, reflecting axonemal tip dynamics observed in cells [50]. Although functions and mechanisms of CCDC66 in cilium assembly are well-characterized in human retinal pigmental epithelial (RPE1) cells, its potential roles in different ciliogenesis pathways and other stages of the cilium cycle, specifically in the maintenance and disassembly of the cilium, remain unknown.

Here, we used localization, proximity mapping and loss-of-function experiments to define the ciliary functions and mechanisms of Ccdc66 in mouse inner medullary collecting duct (IMCD3) cells. We chose IMCD3 cells because they respond to Hedgehog signaling, form epithelial spheroids in three-dimensional matrices, and ciliate via the extracellular pathway, whereas RPE1 cells use the intracellular pathway. We discovered that Ccdc66 depletion disrupts cilium disassembly, length and stability, without affecting assembly in these cells. Ccdc66-depleted cilia exhibited frequent fluctuations in cilium length, increased vesicle and fragment ectocytosis from the cilium tip and were impaired in responding to Hedgehog and Wnt pathway activation. Phenotypic rescue experiments, temporal proximity mapping of Ccdc66 during cilium disassembly and *in vitro* microtubule stabilization assays collectively showed that Ccdc66 regulates these processes by acting as a microtubule-stabilizing factor. Finally, the loss of Ccdc66 disrupted epithelial cell organization and polarity in two-dimensional and three-dimensional cultures. Overall, our results identify Ccdc66 as a regulator of cilium homeostasis, elucidate its role in cellular signaling and tissue organization and provide insight into pathologies linked to Ccdc66.

## Results

### CCDC66 localizes to the ciliary axoneme and regulates cilium length and stability

In RPE1::GFP-CCDC66 stable cells, GFP-CCDC66 localizes to the axoneme and the tip of the primary cilium [41, 43]. In previous studies, we could not detect its ciliary pool in RPE1 cells with the available antibodies. To examine the endogenous localization of Ccdc66 in IMCD3 cells, we generated a custom antibody targeting the 1–756 amino acid (aa) fragment of mouse Ccdc66, which comprises of 935 aa in isoform 1. Next, we stained ciliated IMCD3 cells with antibodies against Ccdc66 and proteins that mark axonemal microtubules (acetylated tubulin) and ciliary membrane (Arl13b) (Fig. 1A). For nanoscale mapping of Ccdc66 localization at the primary cilium, we analyzed its localization using Structured Illumination Microscopy (SIM) and Ultrastructural Expansion Microscopy (U-ExM). Both imaging approaches showed that Ccdc66 co-localized with acetylated tubulin at the axonemal microtubules and basal body (Fig. 1A). Ccdc66 was heterogeneously distributed along the ciliary axoneme in a punctate pattern, similar to the ciliary localization of GFP-CCDC66 in RPE1 cells (Fig. 1A). However, we did not observe a prevalent ciliary tip pool with endogenous staining [43]. These results confirm endogenous localization of Ccdc66 to the ciliary axoneme in mouse cells.

**Figure 1.**
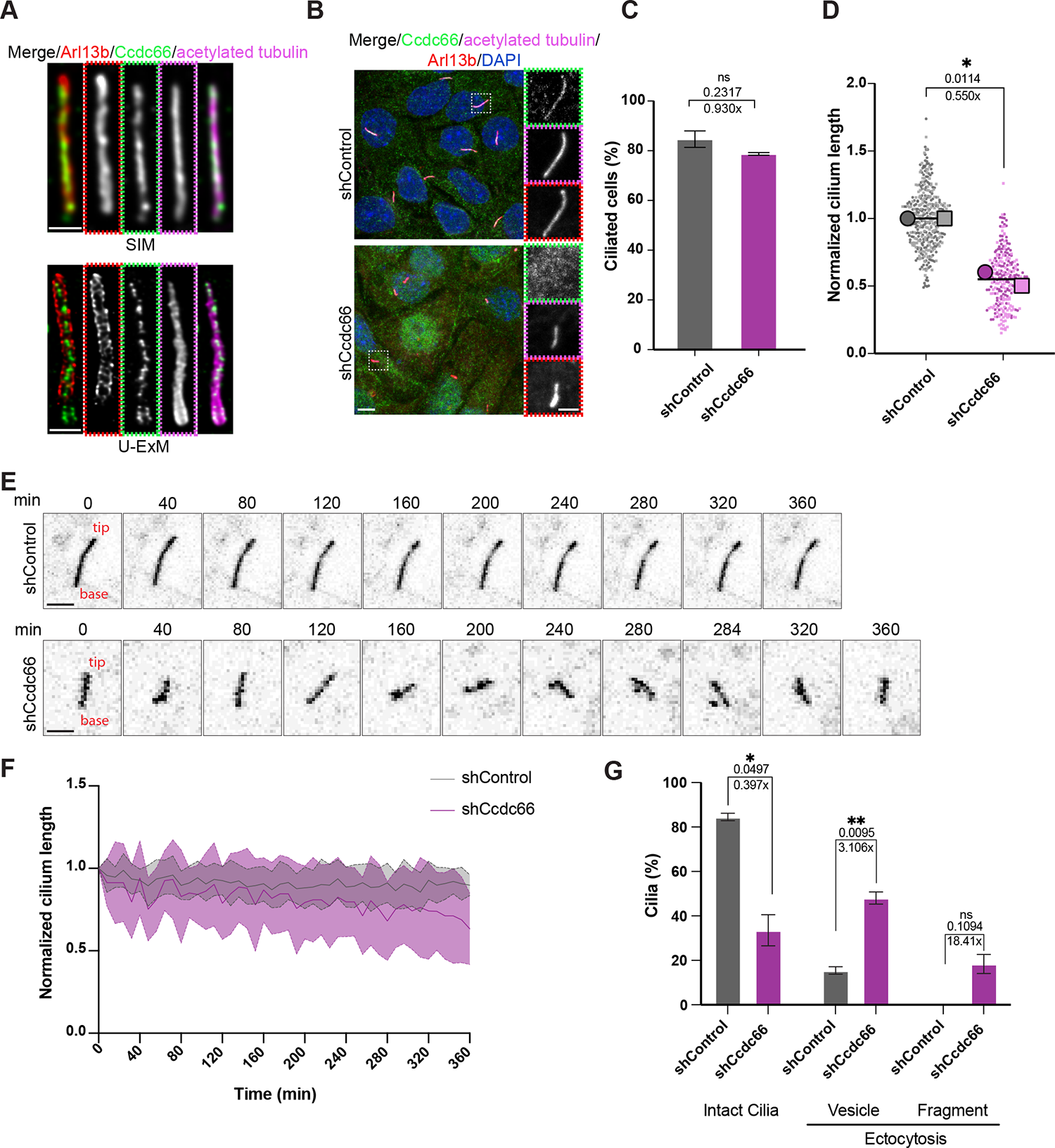
Ccdc66 localizes to the axoneme and regulates primary cilium stability but not cilia formation in IMCD3 cells. (A) Ccdc66 localizes to the axoneme of the primary cilium. Endogenous staining of Ccdc66 in ciliated IMCD3 cells fixed with 4% PFA was performed using homemade antibody raised against the mouse Ccdc66 protein. Cells were co-stained with anti-acetylated-tubulin, anti-Arl13b and DAPI. The top panel shows images obtained using structured illumination microscopy (SIM). The bottom panel shows confocal microscopy images of samples processed by ultra-structure expansion microscopy (U-ExM). Scale bars SIM: 1 µm, Scale bars U-ExM: 222 nm (B-D) Effects of CCDC66 depletion on cilia formation and length. IMCD3 cells transduced and stably expressing either control or shRNA targeting C-terminus of Ccdc66 (shCcdc66) were grown on glass coverslips, fixed with 4% PFA after 48 h of serum starvation and imaged with confocal microscopy. Scale bar: 5 µm. Insets show 3× magnifications of the cilia, Scale bar: 2 µm (C) Quantification of the percentage of cells with cilia (B). Data represents the mean ± SD of 2 independent experiments. n > 600 cells for control and >400 cells for Ccdc66 depletion. The mean cilia percentage is 84.67% for shControl and 78.72% for shCcdc66. (Welch’s t test, ns: not significant) (D) Quantification of cilia length in 3D, according to acetylated-tubulin signal (co-localizing with Arl13b) (B). The super plot represents the mean ± SEM of 2 independent experiments superimposed onto a scatter plot of normalized individual experimental values to control. The first experimental replicate is shown as a darker circle, and the second as a lighter-colored square. Individual values are represented in lighter shades than the corresponding averages. n > 200 cells for each condition. The mean cilia length in Ccdc66-depleted cells decreased to 0.55-fold compared to the mean control length. (Unpaired t test, *p=0.0114) (E-G) Ccdc66 depletion leads to frequent length fluctuations and enhanced ectocytosis in steady-state cilia. (E) The IMCD3::SSTR3-GFP cells transduced with either control or Ccdc66 shRNA virus were grown in FluoroDish and serum starved for 48 h, then imaged with confocal microscopy with 63× objective. Images were acquired every 4 min for 6 h. Still images are inverted to emphasize cilia better. Scale bar: 3 μm. (F) Ciliary kinetics are presented as normalized length curves, measured from SSTR3-GFP fluorescence in two independent experiments, represented as the mean ± SD. n=20 cilia for both shControl and shCcdc66 conditions. (G) Quantification of ciliary ectocytosis events in (E). Ciliary vesicle or fragment ectocytosis were categorized based on the size of the released EVs, vesicle <500 nm and fragment >500 nm, and plotted as a bar plot. (p values of Welch’s t test, *p=0.0497, **p=0.0095, ns=not significant) Ccdc66, coiled-coil domain-containing protein 66; IMCD3, inner medullary collecting duct cell line; PFA, paraformaldehyde; Arl13b, ADP-ribosylation factor-like protein 13B; DAPI, 4′,6-diamidino-2-phenylindole; shRNA, short hairpin RNA; SSTR3, Somatostatin receptor type 3; GFP, green fluorescent protein; EV, extracellular vesicle; MT, microtubules; DMSO, dimethyl sulfoxide; SEM, standard error of mean; SD, standard deviation.

The functions of CCDC66 during cilium assembly and maintenance in IMCD3 cells, which ciliate using the extracellular pathway, remain unknown. To address this, we depleted Ccdc66 from IMCD3 cells by lentivirus-mediated short hairpin RNA (shRNA) treatment and investigated its impact on cilium biogenesis using imaging-based assays. Immunofluorescence, immunoblotting and quantitative PCR experiments confirmed that Ccdc66 was efficiently depleted in IMCD3 cells 6 days after transduction (Fig. S1A-C). The loss of Ccdc66 ciliary and previously reported midbody signal in Ccdc66 shRNA-transduced cells relative to control cells validates the specificity of the rat Ccdc66 antibody (Fig. S1A) [40]. We then examined the effect of Ccdc66 depletion on cilium assembly and length by staining cells for acetylated tubulin and Arl13b. The percentage of ciliated Ccdc66-depleted cells was comparable to that in control cells 48 h post serum starvation (Fig. 1B, 1C).

Additionally, the centrosomal levels of Cep164 and Talpid3 were similar, and Cp110 was removed with comparable efficiency in CCDC66-depleted and control cells (Fig. S1D-F). The lack of defects in the recruitment of distal centriole end proteins Cep164 and Talpid3 required for initiation of ciliation and the removal of the centriole distal cap as marked by Cp110 upon CCDC66 depletion supports the lack of a ciliation defect in IMCD3 cells [51–53]. Despite no apparent ciliation defect, Ccdc66-depleted cells formed shorter cilia with a wide range of lengths, suggesting potential ciliary instability. (Fig. 1B, 2D, S1G). These results show that Ccdc66 is required for the elongation of the axoneme, but not for the initiation of cilium assembly in mouse epithelial cells.

**Figure 2.**
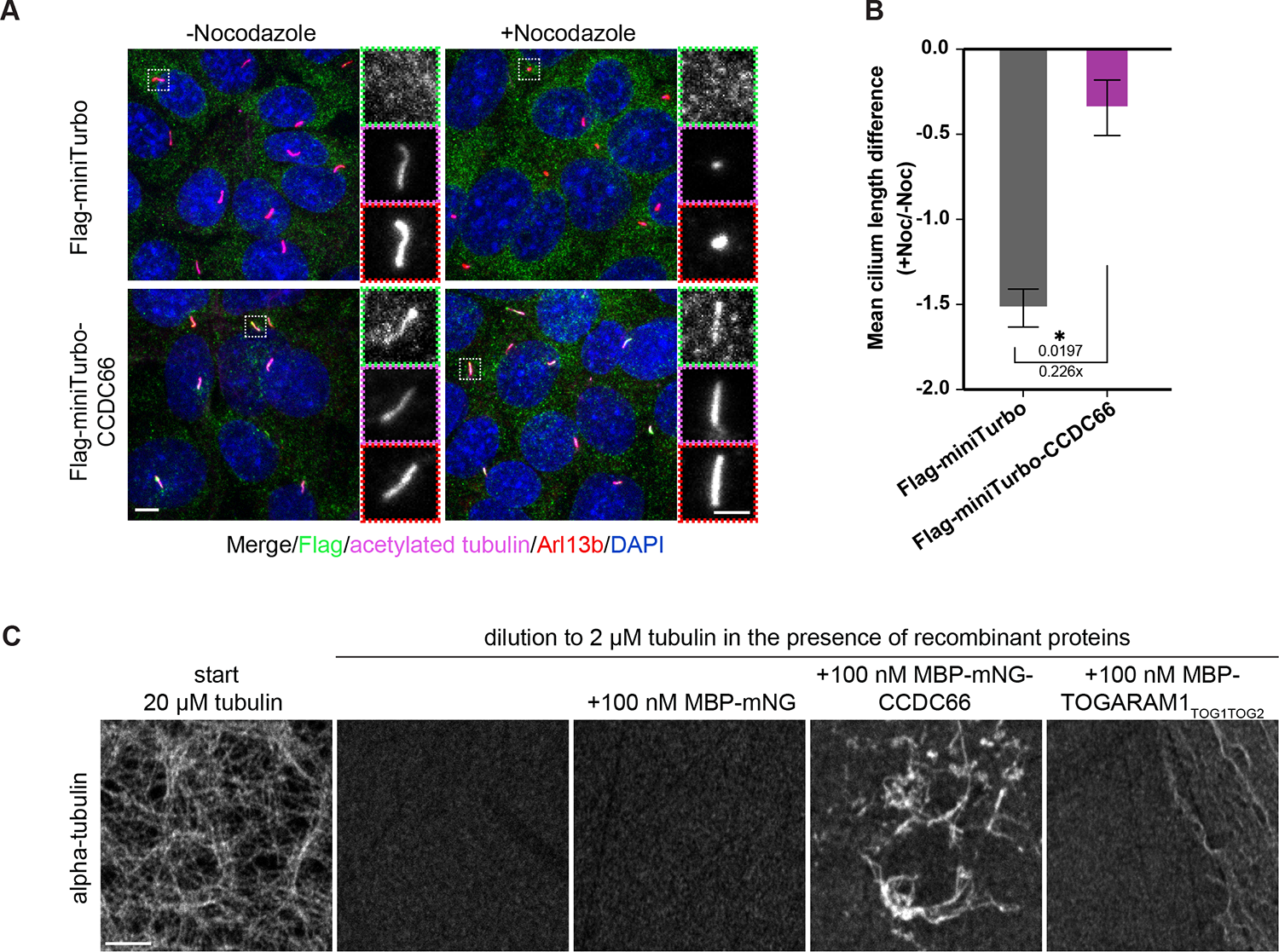
Ccdc66 stabilizes microtubules in cells and *in vitro*. (A, B) IMCD3::miniTurboID-Flag and IMCD3::miniTurboID-Flag-CCDC66 were seeded on coverslips. After 48 h serum starvation, the cells were treated with either DMSO or 400 ng/mL (1.33 µM) Nocodazole for 2h, which destabilizes cytoplasmic and axonemal MTs, fixed with 4% PFA and stained with anti-Flag, anti-acetylated-tubulin, anti-Arl13b and DAPI. Scale bar: 5 µm. Insets show 3× magnifications of the cilia, Scale bar: 2 µm (B) Quantification of average cilia length of +Noc and -Noc treatments relative to their starting length from (A). Data represents mean ± SD of 2 independent experiments. n > 50 cilia for each cell line and treatment per experiment. The mean cilia length difference is −1.521 for miniTurbo control and −0.344 for miniTurbo-CCDC66. (p value of Welch’s t test, *p=0.0197) (C) *In vitro* MT stabilization experiments were performed with MBP-mNeonGreen-CCDC66, MBP-mNeonGreen and MBP-TOGARAM1TOG1TOG2. Fluorescent tubulin mixture (20 μM) was polymerized in BRB80 buffer, 1mM DTT reducing agent, 1 mM GTP and 25% glycerol at 37°C for 30 min. Subsequently, microtubules were diluted to 2 μM tubulin concentration in the buffer without GTP and glycerol, absence or presence of recombinant proteins at indicated concentrations. The reaction was incubated for 5 min at RT, followed by fixation with glutaraldehyde, pelleting onto glass coverslips and additional staining with anti-tubulin antibody. Reactions were imaged using a confocal microscope. Ccdc66, coiled-coil domain-containing protein 66; IMCD3, inner medullary collecting duct cell line; shRNA, short hairpin RNA; DMSO, dimethyl sulfoxide; PFA, paraformaldehyde; Arl13b, ADP-ribosylation factor-like protein 13B; DAPI, 4′,6-diamidino-2-phenylindole; SD, standard deviation; MBP, Maltose-binding protein; BRB80, Brinkley Renaturing Buffer 80; DTT, Dithiothreitol; GTP, Guanosine triphosphate; RT, room temperature.

We investigated the role of Ccdc66 during primary cilium maintenance by analyzing its dynamics, stability and length of steady-state cilia. To measure these ciliary properties spatiotemporally, we performed live imaging of ciliated control and Ccdc66-depleted IMCD3 cells stably expressing the GFP fusion of the ciliary transmembrane protein SSTR3 (GFP-SSTR3). Following 48 h of serum starvation, we imaged cells every 4 min over 6 h with full confocal stacks and quantified cilium length from 3D constructions of the movies (Fig. 1E, 1F). Cilia facing the apical side were analyzed to minimize artifacts that could arise from deformations of basally-oriented cilia due to their interactions with the culture substrate. While cilium length remained relatively constant in control cells, we observed significant variations in length among Ccdc66-depleted cells, supporting defects in cilium stability (Fig. 1F, S1H). Since tubulin PTMs contribute to maintaining cilium stability and length, we quantified the ciliary concentration of polyglutamylated and acetylated tubulin [6]. We found no significant differences between control and CCDC66-depleted cells (Fig. sS1I). This result indicate that cilium instability defects are not due to defective modifications of axonemal tubulins.

Ectocytosis, a process by which vesicles are shed from the cilium, plays a role in controlling ciliary length and maintaining ciliary stability [23, 54, 55]. To test whether ciliary ectocytosis is affected upon CCDC66 loss, we quantified the percentage of ectocytosis events using live-imaging videos. Our analysis showed that the cilia in Ccdc66-depleted cells frequently exhibited two different types of ectocytosis events:

1. vesicle release at the distal end of the cilium, hereafter referred to as vesicle, and
2. breakage of larger ciliary fragments longer than 500 nm, hereafter referred to as fragment. About 66.5% of the cilia observed in CCDC66-depleted cells underwent vesicle and fragment ectocytosis during the imaging period, in contrast to less than 15.5% in control cells (Fig. 1E, 1G). Notably, cilia in Ccdc66-depleted cells showed an increase in the frequency of ectocytosis events. These results suggest that CCDC66 depletion causes fluctuations in cilium length and high incidence of ectocytosis in steady-state cilia by promoting cilium instability.

### CCDC66 regulates primary cilium stability by acting as a microtubule-stabilizing factor

Loss-of-function experiments indicate that Ccdc66 is required for the stability of steady-state cilia. Since CCDC66 is a MAP, we hypothesized that it might regulate cilium stability by acting as a microtubule-stabilizing factor. To test this, we first examined whether CCDC66 overexpression enhances cilia stability by assessing their resistance to microtubule depolymerization in control and CCDC66-expressing cells. Previous studies showed that high doses of nocodazole result in shortened cilia and increased cilium instability [19, 56]. In our experiments, we used previously described IMCD3 cells that stably express FLAG-miniTurbo and FLAG-miniTurbo-CCDC66 [42]. We treated these cells with 400 ng/ml nocodazole for 2 h to induce microtubule depolymerization and challenge axonemal integrity. We then measured the ciliary length difference between cells treated with vehicle control or nocodazole. In CCDC66-expressing cells, the cilium length after nocodazole treatment was 85.3% of its starting length, compared to 41.8% in control cells (Fig. 2A, 2B). These results suggest that cilia in CCDC66-expressing cells are more stable than those in control cells.

CCDC66 promotes microtubule bundling *in vitro*, suggesting that it may stabilize microtubules by crosslinking them [40]. To directly examine the role of CCDC66 in microtubule stabilization, we performed *in vitro* experiments using MBP-mNeonGreen-CCDC66, which was expressed and purified from insect cells as described previously [40]. MBP-mNeonGreen, which does not bind to microtubules, and the first two TOG domains of TOGARAM1, which bind free tubulin but not the microtubule lattice and promote microtubule polymerization but not stability *in vitro*, were used as controls [50, 57]. We polymerized rhodamine-labeled microtubules from 2 µM tubulin mixture in the presence of GTP and glycerol at 37°C for 30 min. These microtubules were then diluted 10x in prewarmed buffer, either with and without 100nM MBP-mNeonGreen-CCDC66, MBP-mNeonGreen, or MBP-TOGARAM1_TOG1TOG2_. The mixtures were incubated at room temperature for 5 min. While microtubules disassembled in the presence of MBP-mNeonGreen and MBP-TOGARAM1_TOG1TOG2_, microtubule bundles induced by MBP-mNeonGreen-CCDC66 persisted after dilution (Fig. 2B). These results show that CCDC66 stabilizes microtubules against dilution-induced depolymerization. Our loss-of-function, overexpression, and *in vitro* experiments suggest that CCDC66 exerts its ciliary functions via stabilizing axonemal microtubules.

### CCDC66 depletion enhances cilium disassembly by regulating AURKA and HDAC6

Maintenance of cilium length requires a dynamic balance between cilium assembly and disassembly processes [22, 58, 59]. To determine whether and how Ccdc66 contributes to this process, we quantified the percentage of ciliated cells upon induction of cilium disassembly using serum stimulation. To this end, IMCD3 cells were infected with control and Ccdc66-targeting lentivirus and then subjected to serum starvation for 48 h to induce cilium assembly. Subsequently, we quantified the percentage of ciliated cells over a 12 h serum stimulation time course (Fig. 3A).

**Figure 3.**
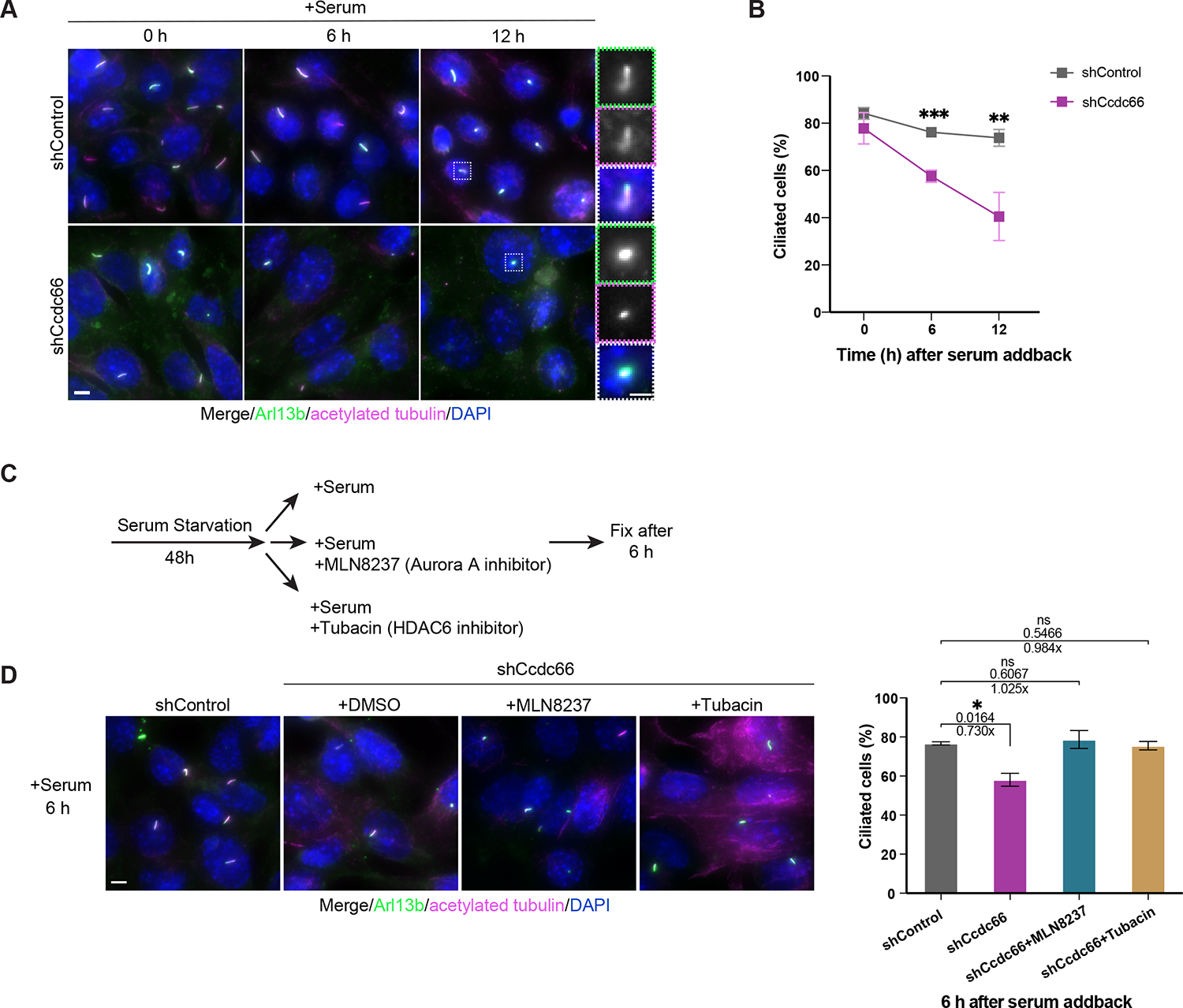
Cilia disassemble more rapidly in Ccdc66-depleted cells, which is regulated by the AurkA-HDAC6 axis. (A, B) Ccdc66 depletion leads to enhanced disassembly. Control and Ccdc66-depleted cells were serum starved for 48 h, then serum stimulated for either 6h or 12h in total. Cells were fixed at indicated time points with 4% PFA, stained against anti-acetylated-tubulin, Arl13b and DAPI, and imaged with widefield fluorescence microscopy. Scale bar: 5 µm. Insets show 3× magnifications of the cilia, Scale bar: 2 µm (B) Quantification of ciliated cell percentagein (A). Data represents mean ± SD of 3 independent experiments. n > 600 cells for control and >400 cells for Ccdc66 depletion per experimental replicate. Mean cilia percentage at 0 h is 84.16% for shControl, 77.84% for shCcdc66; at 6 h is 76.15% for shControl, 57.56% for shCcdc66; at 12 h is 73.83% for shControl, 40.51% for shCcdc66. (p values of multiple t test of grouped data, ns not significant, ***p=0.00032, **p=0.0059). (C, D) AURKA and HDAC6 inhibition rescues enhanced cilium disassembly phenotype of Ccdc66-depleted cells. (C) The experimental plan for rescue of cilia disassembly phenotype through inhibition of AurkA and HDCA6: control and Ccdc66 shRNA-depleted cells were serum starved for 48 h and released into serum-rich media to induce cilia disassembly for 6 h with DMSO (control), 500 nM AurkA inhibitor, MLN8237, or 2 µM HDAC6 inhibitor, Tubacin. Following incubation with inhibitors, cells were fixed and stained for acetylated-tubulin, Arl13b and DAPI. (D) Widefield microscopy of treated cells as in (C) with representation of Ccdc66-depleted and treated cells compared to control untreated cells, with quantification of cilia number. Data represents mean ± SD of 2 independent experiments. n > 700 cells for control, n > 400 cells for CCDC66 depletion, n > 600 for CCDC66 depleted cells with inhibitors per experimental replicate. The mean cilia percentage at 6 h is 76.74% for shControl, 58.11% for shCcdc66, 78.73% for shCcdc66+MLN8237 and 75.56% for shCcdc66+Tubacin. (p values of one-way ANOVA with Tukey’s post hoc test *p=0.0164, ns: not significant) Ccdc66, coiled-coil domain-containing protein 66; PFA, paraformaldehyde; SD, standard deviation; AurkA, Aurora kinase A; HDAC6, Histone deacetylase 6.

Notably, there was a rapid and significantly greater decrease in the percentage of ciliated cells upon CCDC66 depletion as compared to control cells at both 6 and 12 h after serum stimulation (Fig. 3A, 3B). In control cells, the percentage of ciliated cells decreased from 84.16% ± 2.5% to 73.83% ± 3.5% over 12 h. However, in CCDC66-depleted cells, the percentage significantly decreased to 40.51% ± 10.2% within the same period (Fig. 3B). Strikingly, majority of the Ccdc66-depleted cells had ciliary stubs, identified by a short (<1 μm) cilium stained positive for both acetylated tubulin and Arl13b (Fig. 3A). The enhanced deciliation phenotype upon CCDC66 depletion suggests that it acts as an inhibitor of cilium disassembly, potentially by stabilizing the axoneme.

To validate the specificity of the ciliary phenotypes in mouse cells, we performed phenotypic rescue experiments with human CCDC66 shRNA-resistant protein. For these experiments, we used IMCD3 cells stably expressing Flag-miniTurbo-CCDC66 (hereafter miniTurbo-CCDC66) [42]. In both control and Ccdc66-depleted ciliated cells, miniTurbo-CCDC66 localized to the basal body, axoneme, ciliary tip and centriolar satellites (Fig. S2A). Notably, the satellite pool of miniTurbo-CCDC66 became more prominent throughout the 12 h of serum stimulation. miniTurbo-CCDC66 expression rescued both the enhanced cilium disassembly and the shorter cilium length phenotype in CCDC66-depleted cells, restoring them to levels comparable to those in control cells (Fig. S2A-C). This demonstrates that these phenotypes are specific to Ccdc66 depletion.

We previously showed that CCDC66 interacts with the ciliary MAP CEP104 and cooperates with it to regulate cilium length in RPE1 cells [42]. Therefore, we investigated the functional dependency between CCDC66 and CEP104 during cilium disassembly and axoneme elongation. To this end, we examined whether stable expression of CEP104-BirA*-FLAG (hereafter CEP104-BirA*) restores cilium disassembly and length defects associated with Ccdc66 depletion. In both control and Ccdc66-depleted ciliated cells, CEP104-BirA* localized to the basal body and the ciliary tip, with axonemal localization also observed in some cells (Fig. S2D). In CEP104-BirA*-expressing stable cells, Ccdc66 depletion resulted in a rapid and significantly greater decrease in the percentage of ciliated cells as compared to control cells at both 6 and 12 h after serum stimulation (Fig. S2D, S2E). Similarly, CEP104-BirA expression did not compensate for the reduced cilium length defect observed in CCDC66-depleted cells (Fig. S2F). These findings suggest that, unlike in RPE1 cells, Ccdc66 does not cooperate with Cep104 to regulate cilium length and disassembly in IMCD3 cells. Instead, Ccdc66 might act upstream of Cep104 by recruiting it to the ciliary axoneme. This variation could also be due to differences in species and tissue origin, ciliogenesis pathways, and cellular transformation processes.

A major mechanism underlying cilium disassembly involves the phosphorylation of histone deacetylase 6 (HDAC6) by Aurora A kinase (AurkA) and subsequent deacetylation of axonemal tubulins [8, 33, 60]. To investigate if Ccdc66 plays a role in regulating cilium disassembly through a pathway dependent on HDAC6 and AurkA, we performed cilium disassembly experiments in control and Ccdc66-depleted cells treated with tubacin and MLN as specific inhibitors to target HDAC6 deacetylase and AurkA kinase activity, respectively (Fig. 3C). Treatment with both tubacin and MLN effectively rescued the enhanced cilium disassembly phenotype observed in Ccdc66-depleted cells at 6 h (Fig. 3D). These findings show that CCDC66 regulates cilium disassembly by acting via the disassembly pathway involving HDAC6 and AURKA, possibly downstream of their decision-to-initiate-cilia loss activity [37].

### CCDC66 depletion results in cilium length fluctuations and increased ectocytosis during disassembly

Events underlying cilium disassembly was reported to vary within a single-cell population [37]. In IMCD3 cells, these events were characterized as rapid deciliation through whole-cilium shedding (instant loss), gradual resorption or a combination of both, starting with gradual resorption followed by rapid deciliation. To examine the role of CCDC66 in the kinetics of cilium disassembly and individual cilium disassembly events, we monitored spatiotemporal dynamics of cilium disassembly in IMCD3::GFP-SSTR3 cells. These cells were infected with control and Ccdc66-targeting virus, serum starved for 48 h, induced with serum stimulation and imaged using confocal microscopy, capturing full stack images every 4 min over 6 h after serum addition (Fig. 4A). From 3D constructions of these movies, we quantified cilium length in cells with disassembling cilia. While the decrease in cilium length in control cells followed a largely linear pattern, CCDC66-depleted cells displayed significant variations in length (Fig. 4A, 4B). Notably, cilia in control cells disassembled largely through gradual resorption. However, upon Ccdc66 depletion, we observed an increase in the frequency of cilia disassembling either through instant loss or a combination of gradual resorption and instant loss (Fig. 4A, S3A). These findings support the role of CCDC66 in maintaining cilium stability.

**Figure 4.**
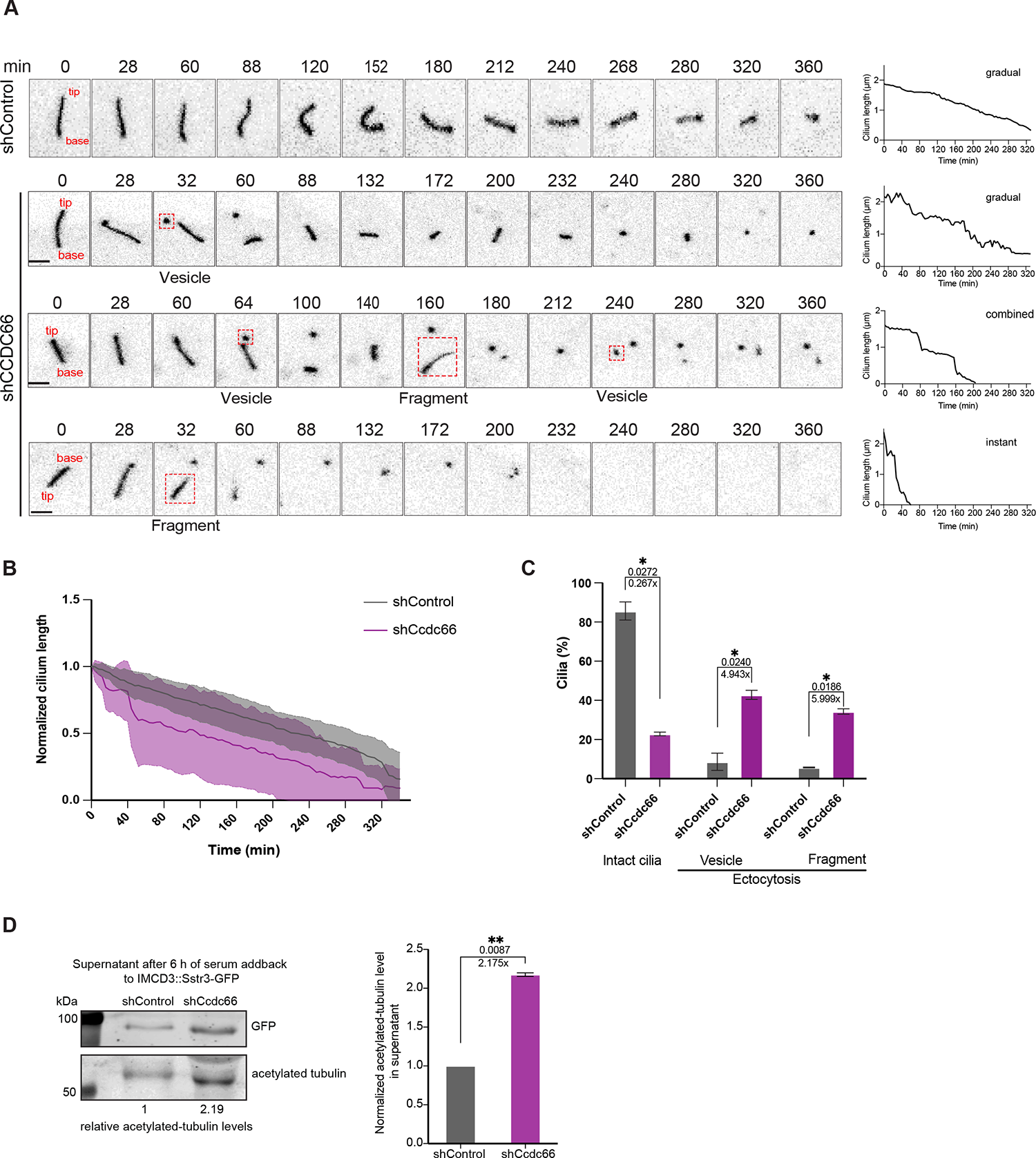
Live imaging of Ccdc66-depleted cells reveals an increase in ectocytosis and frequent fluctuations in cilia length. (A-C) Cilium disassembly kinetics upon serum stimulation. The IMCD3::SSTR3-GFP cells transduced with either control or Ccdc66 shRNA were grown and serum starved in FluoroDish for 48 h, then imaged with confocal microscopy for 6 h immediately upon serum addition. With images acquired every 4 min. Representative images show one control cilium that gradually resorbs, and three Ccdc66-depleted cilia that undergo disassembly through gradual resorption, instant loss (whole cilium shedding), or a combination of both. Still images are inverted to emphasize cilia better. Scale bar 3 μm (B) Normalized cilia length curves are measured from fluorescence of two independent experiments and represented as the mean ± SD. n=20 cilia for both shControl and shCcdc66 conditions. (C) Quantification of ciliary ectocytosis events in (A). Ciliary events, including vesicle releases (<500 nm) and cilium fragment shedding (>500 nm), were categorized based on the size of the released extracellular vesicles (EVs) and plotted as a bar plot (p values of Welch’s t test *p=0.0272, *p=0.0240, *p=0.0186). (D) Media from IMCD3::SSTR3-GFP was collected 6 h post serum stimulation and separated by high speed centrifugation for subsequent western blot. Pelleted material shows the presence of ciliary markers at expected molecular weights suggesting pelleting of ciliary EVs and fragments. Membranes are blotted with anti-GFP and anti-acetylated tubulin. Shown in the graph is the fold change in the intensity of the acetylated tubulin bands, normalized to shControl of the displayed experimental replicate. The sample was normalized to total cell protein abundance before loading to account for differences in cell numbers between control and Ccdc66-depleted cells. n=2. The bar plot represents mean ± SD of 2 independent experiments. (One sample t test p value **p=0.0087) Ccdc66, coiled-coil domain-containing protein 66; IMCD3, inner medullary collecting duct cell line; SSTR3, Somatostatin receptor type 3; GFP, green fluorescent protein; shRNA, short hairpin RNA; SD, standard deviation.

To investigate the effect of CCDC66 depletion on ciliary ectocytosis during cilium disassembly, we used live imaging videos to quantify the percentage of different ectocytosis events. We found that about 77.12% of the cilia in CCDC66-depleted cells underwent ectocytosis over 6 h of imaging, compared to less than 14.4% in control cells (Fig. 4A, 4C). In a complementary analysis, we purified extracellular vesicles (EVs) from the supernatants of serum-starved control and Ccdc66-depleted IMCD3::GFP-SSTR3 cells 6 h after serum stimulation.

Immunoblotting against GFP-SSTR3 and acetylated tubulin revealed a higher concentration of ciliary proteins in the EVs from Ccdc66-depleted cells compared to control cells (Fig. 4D). Given that EVs can be released by both ciliated and non-ciliated cells, the detection of ciliary proteins within these vesicles implies that they predominantly originate from cilia. Collectively, these findings further support that loss of CCDC66 enhances cilium disassembly, resulting in fluctuations in cilium length and increased ectocytosis by promoting cilium instability.

### Temporal proximity mapping of CCDC66 during cilium disassembly reveals new regulators of cilium length and stability

To gain insight into how CCDC66 regulates cilium length, stability and disassembly, we identified its proximity interaction partners during cilium disassembly in IMCD3 cells. To benchmark our data against proximity cilium proteomes and to temporally probe the proximity partners of CCDC66 during cilium disassembly, we used IMCD3 cells that stably express miniTurbo-CCDC66 [42, 61, 62]. For proximity mapping experiments, we serum-starved miniTurbo or miniTurbo-CCDC66-expressing cells for 48 h and then induced cilium disassembly and biotinylation using serum and biotin stimulation. Subsequently, we validated the localization and induced biotinylation of miniTurbo-Flag-CCDC66 by staining cells for FLAG to mark the fusion protein, streptavidin to mark the biotinylated proteins, either ARL13B or acetylated tubulin to mark the cilium and/or PCM1 to mark the centriolar satellites (Fig. 5A, 5B). As assessed by immunofluorescence analysis of cells induced by 1 and 6 h serum stimulation and a 30 min biotinylation, miniTurbo-CCDC66 localized and biotinylated proteins at the centriolar satellites, basal body, and cilium in serum-starved cells (Fig. 5A). Consistent with previous reports, centriolar satellite pool of CCDC66 decreased in ciliated cells and increased upon cilium disassembly (Fig. 5B) [42].

**Figure 5.**
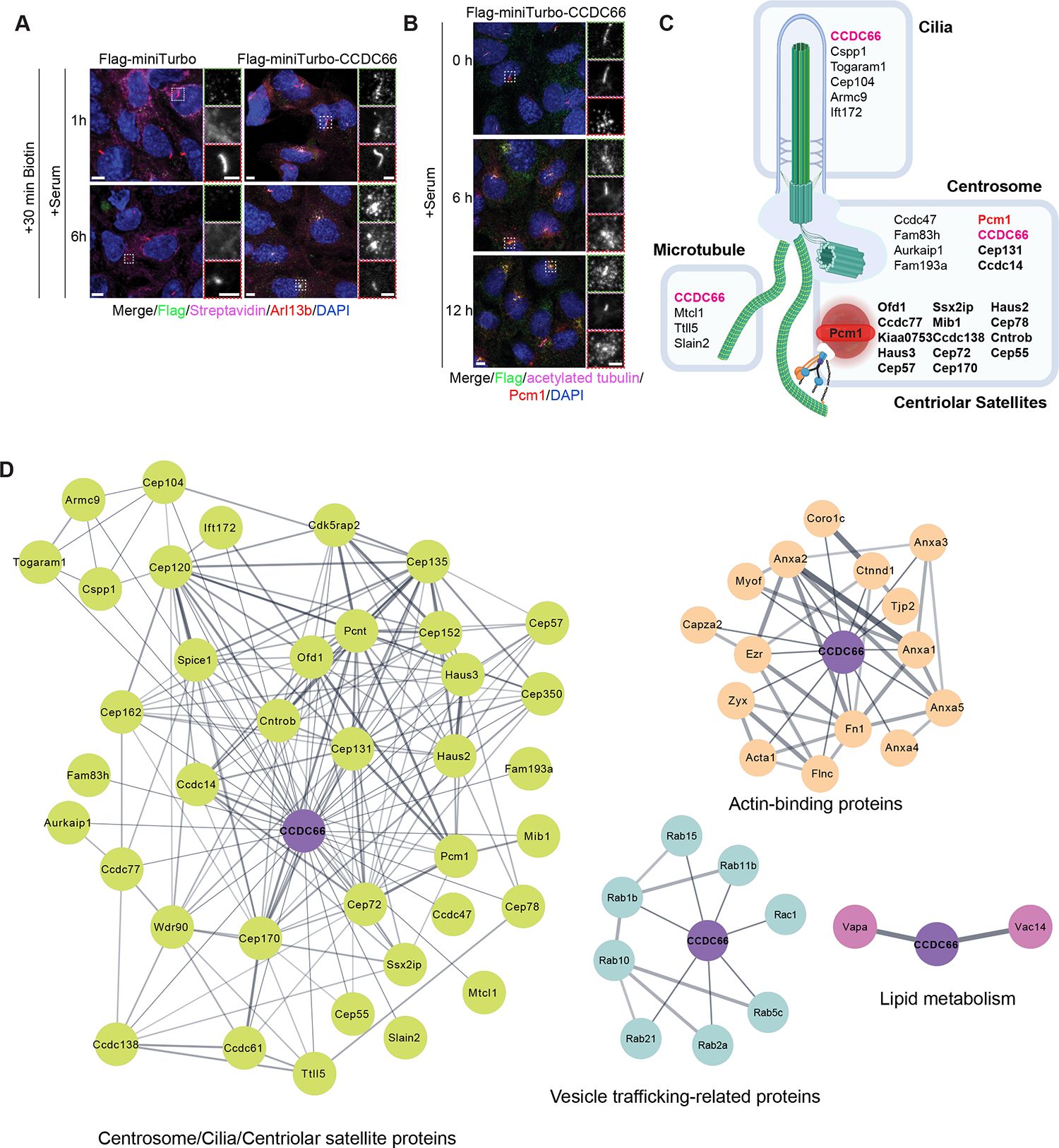
Temporal proximity interactomes of Ccdc66 during cilium disassembly (A) The ciliated Flag-miniTurboID or Flag-miniTurboID-CCDC66 IMCD3 cell lines were seeded on glass coverslips and treated with 500 μM biotin within the last 30 min of a 1 h or 6 h serum stimulation. Cells were then fixed with methanol and stained with anti-Flag, Streptavidin-Alexa568 coupled, anti-Arl13b, and DAPI, and imaged with confocal microscopy. Scale bar: 5 µm. Insets show 3× magnifications of the cilia, Scale bar: 2 µm (B) Ccdc66 translocates to centriolar satellites upon serum stimulation and cilia disassembly. IMCD3::Flag-miniTurbo-CCDC66 were fixed with 4% PFA at the indicated time points of serum stimulation and stained with anti-Flag, anti-PCM1, anti-acetylated-tubulin, and DAPI. Scale bar: 5 µm. Insets show 3× magnifications of the cilia, Scale bar: 2 µm (C) Schematic diagram of sub-interaction networks of Ccdc66 interactome at 1 h and 6 h disassembly conditions based on cellular compartment. The Ccdc66 proximity interactors were grouped using DAVID functional annotation tool and literature mining. The interaction networks visualized for the cellular compartments of the centrosome, cilia, and centriolar satellite were categorized to represent their respective unique compartments. Proteins highlighted in bold are the ones that localize to both centrosomes and centriolar satellites. (D) The sub-interaction networks of the Ccdc66 interactome at 1 h and 6 h disassembly conditions. Ccdc66 proximity interactors were grouped using the DAVID functional annotation tool and literature mining. The interaction networks visualized for for cellular compartments include centrosome/cilia/centriolar satellite and actin cytoskeleton. Those visualized for biological process include vesicular trafficking and lipid metabolism. The interconnectedness among the proteins in each network was determined using the STRING database.

After validation of the cell line, we performed large scale streptavidin pulldown of biotinylated proteins from ciliated cells serum stimulated for 1 and 6 h, analyzed them by mass spectrometry and defined high confidence CCDC66 interactome using Normalized Spectral Abundance Factor (NSAF) analysis [63] (Fig. S4A). This yielded 282 and 301 high-confidence interactors for CCDC66 after 1 hour and 6 hour serum stimulation, respectively (Table 1) (Fig. S4B). Of these, 211 proteins were shared between 1 hour and 6 hour interactomes (Fig. S4B). To comprehensively analyze the CCDC66 proximity interactome during disassembly, we merged the proximity interactors from two time points (1 and 6 hours), collectively referred to as the CCDC66 disassembly interactome. Analysis of this interactome, through Gene Ontology (GO) analysis and literature mining, revealed enrichment of centrosome, cilium, and satellite proteins, MAPs, actin-binding proteins, and proteins involved in vesicle trafficking and lipid metabolism (Fig. 5C, 5D, S4C). We organized the proteins from these categories into a sub-interaction network by combining STRING database and ClusterONE plug-in on Cytoscape, which revealed potential functional clusters and multiprotein complexes (Fig. S5D).

We also cross-referenced the CCDC66 disassembly interactome with the ciliated proximity interactome of CCDC66, both of which were generated using the same cell lines [42]. For the comparative analysis, we reanalyzed the raw ciliated CCDC66 interactome using the same filtering thresholds that were applied to the disassembly interactome, to account for variations in methodologies and stringency levels. This revealed 202 proteins were shared between ciliated and disassembly interactomes of CCDC66 (Fig. S4B). For visualization of the comparative analysis of the ciliated and disassembly interactomes of CCDC66, we plotted log_2_(NSAF) values in heat maps representing proteins linked to centrosome, cilia, centriolar satellites, actin, vesicles, and lipid metabolism (Fig. S4D-G).

Several CCDC66 disassembly proximity interactors stood out due to their known relationship to CCDC66 or their roles in regulating cilium length, stability and disassembly. Consistent with the redistribution of CCDC66 from the primary cilium to centriolar satellites, the CCDC66 disassembly interactome was enriched for a subset of centriolar satellite proteins compared to the ciliated proteome (Fig. 5D, S4D). The interactome also included components of a Joubert syndrome module linked to cilium length regulation, Armc9, Togaram1, Cep104 and Cspp1 (Fig. 5C, S4D) [49, 57, 64]. Considering the role of actin cytoskeleton in cilium length regulation and the presence of actin-binding proteins in the cilium proximity interactome, the depletion or enrichment of actin-binding proteins in the CCDC66 disassembly interactome is also important (Fig. 5D, S4E) [65–67]. Additionally, given the importance of vesicular trafficking in regulating cilium, identification of proteins associated with vesicle trafficking such as Rab and Rab-like membrane trafficking proteins and lipid metabolism is also notable (Fig. 5D, S4F, S4G) [68]. Finally, we analyzed the CCDC66 disassembly proteome for proteins known to be involved in cilium disassembly. Although inhibiting AurkA and HDAC6 rescued CCDC66 disassembly phenotypes, our analysis did not detect these key players in cilium disassembly interactome. This indicates that CCDC66 has an indirect relationship with these proteins. However, the interactome included CEP55 and components of the CCT chaperonin complex, which were shown to regulate cilia disassembly via stabilization of AurkA (Fig. 5, Table 1) [69].

### CCDC66 depletion perturbs cilium content regulation and ciliary signaling

To investigate how ciliary defects associated with CCDC66 loss affect cilium function, we examined the response to Hedgehog pathway activation and Wnt pathway response in Ccdc66-depleted cilia. Upon Hedgehog ligand stimulation, the GPCR SMO enters the cilium while GPR161 exits, ultimately leading to the transcriptional activation of Hedgehog target genes [70]. As functional readouts for Hedgehog pathway activation, we quantified the ciliary entry efficiency of SMO and the upregulation of Gli1 and Ptch1 (Fig 6A-C). To this end, ciliated control and Ccdc66 depleted cells were treated with 200 nM Smoothened agonist (SAG) for 4 h and analyzed by immunofluorescence and quantitative PCR. After SAG stimulation, the ciliary level of SMO decreased significantly in Ccdc66-depleted cells compared to control cells, and the percentage of SMO-positive cilia also decreased (Fig. 6A, 6B). These results show that Ccdc66 loss disrupts SAG-induced ciliary accumulation of SMO. Moreover, we determined the downstream consequences of these alterations by quantifying Gli1 and Ptch1 transcriptional upregulation in SAG-treated cells. As compared to control cells, Ccdc66-depleted cells showed defects in Gli1 and Ptch1 upregulation in response to SAG treatment (Fig. 6C). In addition to Hedgehog signaling, primary cilium has been reported for its functions in Wnt signaling [71, 72]. To assess the role of Ccdc66 in Wnt signaling, we compared basal mRNA expression levels of Wnt effector genes between control and Ccdc66-depleted cells. We tested one of the most responsive downstream target gene of β-catenin, and target genes involved in cell adhesion and cellular metabolism. The Ccdc66-depleted cells exhibited defects in the expression of Axin2, Pdk1, and Fibronectin compared to control cells (Fig. 6D). These findings collectively demonstrate that Ccdc66 is required for both Hedgehog and Wnt signaling responses.

**Figure 6.**
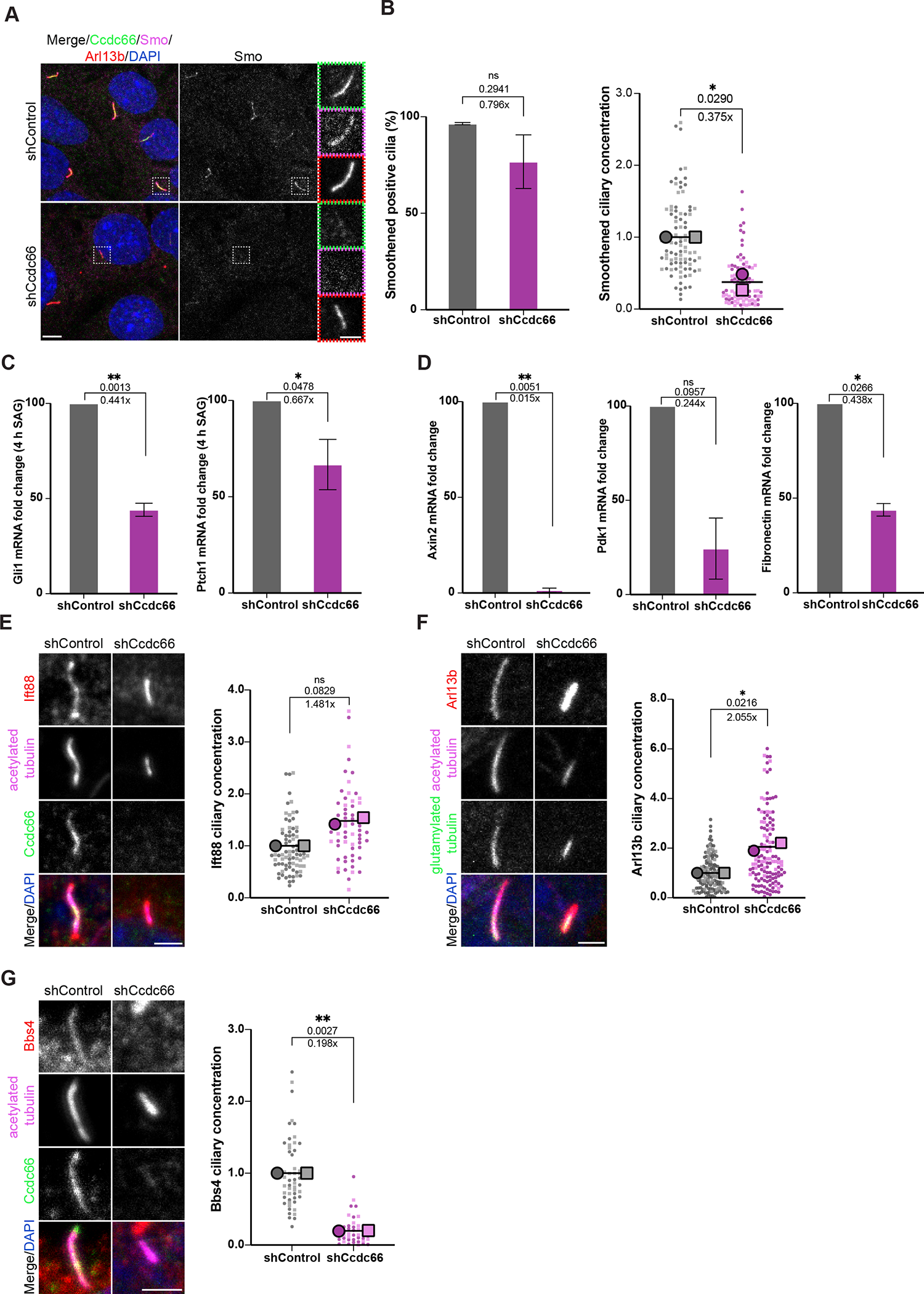
CCDC66-depleted cells exhibit defective ciliary signaling and transport. (A, B) Control and Ccdc66-depleted cells were ciliated for 48h, followed by treatment with 100 nM SAG for 4 h, then fixed with 4% PFA and processed for immunofluorescence with anti-Smoothened, anti-Ccdc66, anti-Arl13b and DAPI. Scale bar: 5 µm. Insets show 3× magnifications of the cilia, Scale bar: 2 µm (B) Quantification of Smo-positive cilia and ciliary Smo concentration in (A). Data represents mean ± SD for smoothened positive cilia and mean ± SEM for ciliary concentration of 2 independent experiments. n =100 cells for control and Ccdc66 depletion. The mean cilia percentage is 96.46% for shControl and the mean cilia percentage is 76.75% for shCcdc66. (p value of Welch’s t test ns: not significant). The mean ciliary fluorescence density of Smo in Ccdc66 depletion is 0.374-fold decreased compared to the control mean. (Welch’s t test p value *p=0.029) (C) Effect of Ccdc66 depletion on Hedgehog pathway activation as assessed by quantitative PCR. mRNA isolated from control and Ccdc66-depleted cells, ciliated for 48 h and treated with SAG for 4 h, was analyzed with primers recognizing Gli1 and Ptch1, and β-actin primers as normalization control. Box plots shows means ± SD. n=3. Below p values presented is fold change. (One sample t test p values **p=0.0013 and *p=0.0478) (D) Effect of CCDC66 depletion on Wnt pathway as assessed by quantitative PCR. mRNA isolated from control and Ccdc66-depleted cells, ciliated for 48 h was analyzed with primers recognizing Axin2, PDK1and Fibronectin, using β-actin primers as normalization control. Box plots shows means ± SD. n=2. Below p values presented is fold change. (One sample t test p values **p=0.0051, *p=0.0296, ns=not significant) (E) 48 h-ciliated control and Ccdc66-depleted cells were fixed with 4% PFA and stained against anti-IFT88, anti-Ccdc66 and anti-acetylated-tubulin. Ift88 signal was measured from Ccdc66-depleted cells based on the absence or visible decrease of Ccdc66 signal. Measured are ciliary integrated densities=mean intensity*area to represent ciliary signals of cilia varying in size. The super plot of normalized individual experimental values with mean± SEM represents 2 independent experiments. The different colored experimental replicates are shown as either circle or squares and with different color lightness. n > 50 cells for each condition. The mean ciliary fluorescence density of Ift88 in CCDC66 depletion is 1.48-fold higher than the control mean. (Welch’s t test p value ns: not significant) (F) Quantification of ciliary Arl13b in control and CCDC66-depleted cells, serum starved and stained against acetylated and polyglutamylated-tubulin and Ar13b to mark cilia. Measured are ciliary integrated densities. Super plot represents normalized individual experimental values and means ± SEM of 2 independent experiments. n > 150 cilia for each condition. (p value of unpaired t tests *p=0.0216) (G) Control and Ccdc66-depleted cells were fixed with 4% PFA and stained against anti-Bbs4, anti-Ccdc66 and anti-acetylated-tubulin. Fluorescence intensity of BBS4 at the cilia were assessed as in (E). The super plot of normalized individual experimental values with mean± SEM represents 2 independent experiments. n > 50 cells for each condition. Mean ciliary fluorescence density of BBS4 in Ccdc66 depletion is 0.198-fold decreased compared to the control mean. (Welch’s t test p value **p=0.0027) Ccdc66, coiled-coil domain-containing protein 66; IMCD3, inner medullary collecting duct cell line; Smo, Protein smoothened; SAG, Smoothened agonist; Arl13b, ADP-ribosylation factor-like protein 13B; DAPI, 4′,6-diamidino-2-phenylindole; Gli1, Zinc finger protein GLI1; Pthc1, Protein patched homolog 1; PDK1, Pyruvate dehydrogenase (acetyl-transferring) kinase isozyme 1; PFA, paraformaldehyde; IFT88, Intraflagellar transport protein 88 homolog; BBS4, Bardet-Biedl syndrome 4 protein; SD, standard deviation; SEM, standard error of mean.

Defective ciliary signaling in Ccdc66-depleted cells may result from impaired intraciliary transport. Ciliary content, and consequently their signaling competence, is established by the intraflagellar transport (IFT) machinery involving IFT-A, IFT-B and BBSome complexes [1, 73]. To investigate the impact of CCDC66 depletion on IFT, we examined the recruitment of the IFT-B machinery to the basal body and cilium.

We observed an increase in Ift88 at the cilium and basal body in Ccdc66-depleted cells, although this increase was not statistically significant (Fig. 6E). Moreover, these cells exhibited increased ciliary levels of the small GTPase Arl13b, which is important for regulation of Hedgehog signaling and IFT (Fig. 6F). Next, we measured the ciliary levels of Bbs4, a component of the BBSome that redistributes between centriolar satellites and the primary cilium, similar to CCDC66. In Ccdc66-depleted cells, the levels of Bbs4 at the cilium were reduced compared to control cells, similar to previous observation in RPE1 cells [41] (Fig. 6G). Together, these findings suggest that CCDC66 depletion disrupts ciliary signaling in part by compromising intraciliary transport.

### CCDC66 depletion perturbs epithelial cell organization

Proper cilium biogenesis and signaling are crucial for epithelial cell organization [74–76]. When grown in a three-dimensional (3D) gel matrix, epithelial cells organize into polarized, spheroid structures that reflect the *in vivo* organization of the epithelial tissues. To investigate the consequences of ciliary defects associated with CCDC66 loss in tissue architecture, we performed CCDC66 loss-of-function experiments using the 3D spheroid cultures of IMCD3 cells, which mimic *in vivo* organization of the kidney collecting duct [74]. To this end, control and Ccdc66-depleted IMCD3 cells were cultured in Matrigel for 4 days, serum-starved for 2 days, and spheroid architecture was visualized by staining cells for markers for cilia (Arl13b), cell–cell contacts and polarity (actin and β-catenin). Control cells formed spheroids characterized by a central lumen, apically oriented cilia and well-organized apical and basal surfaces (Fig. 7A). In contrast, Ccdc66-depleted cells were impaired in their ability to form structurally proper spheroids compared to the control group (Fig. 7B).

**Figure 7.**
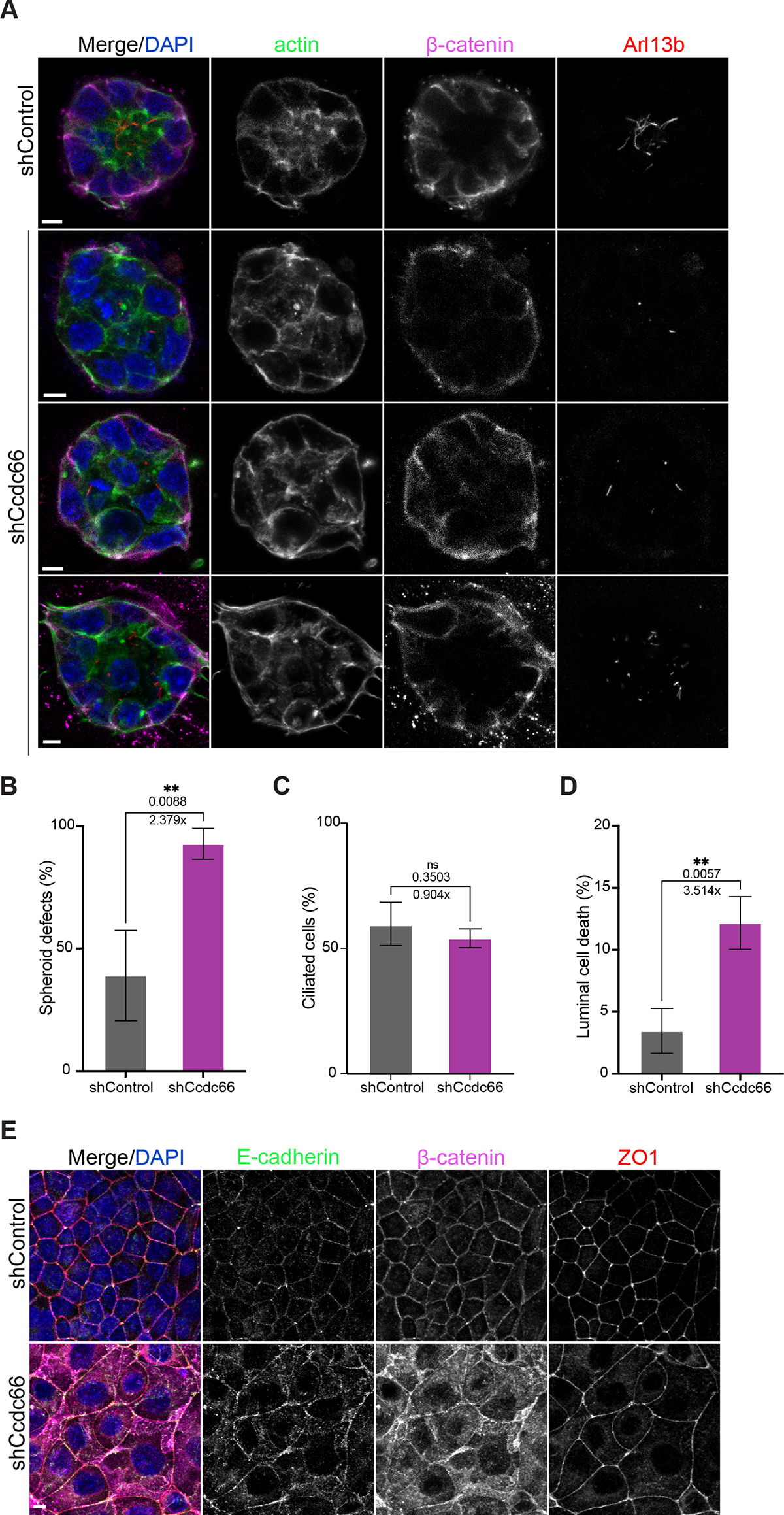
Ccdc66 depletion disrupts tissue organization in 3D and 2D epithelial cultures. (A) IMCD3 cells transduced and stably expressing either control shRNA or Ccdc66 shRNA were grown layered on 100% Matrigel for 4 days before transferring into serum-free media to induce ciliation for 48 h. Cells organized into epithelial spheroids in Matrigel, which were fixed with 4% PFA and processed for immunofluorescence analysis with Phalloidin (staining F-actin), β-catenin and Arl13b, to observe apicobasal polarity and ciliation. Representative images show regular control spheroids and Ccdc66-depleted spheroids with visibly disrupted apicobasal polarity, absence of lumen or cells protruding into lumen, as well as short misoriented cilia. Scale bar: 5 µm. (B-D) Quantification of spheroid defects in (A). n=37 spheroids for control and n=38 spheroids for Ccdc66 depletion. The mean defect percentage is 38.99 for shControl and 92.72 for shCCDC66, with fold change increase of 2.38. (p value of Welch’s t test **p=0.0088). (C) Quantification of cilia number in (A). Data represents mean ± SD of 3 independent experiments. Mean cilia percentage is 59.78 for shControl and 54.02 for shCcdc66, with fold change decrease of 0.90. (p value of Welch’s t test ns: not significant). (D) Quantification of luminal cell death in (A). Percentage of apoptosis was scored based on DAPI and β-catenin staining and distance of apoptotic bodies within spheroid lumen. Data represents mean ± SD of 3 independent experiments. Mean luminal cell death percentage is 3.46 for shControl and 12.16 for shCcdc66, with fold change increase in cell death of 3.51. (p value of Welch’s t test **p=0.0057). (E) Control and Ccdc66-depleted cells were grown to 100% confluency to induce formation of cell-to-cell contacts, then fixed with 4% PFA and stained for tight-junction marker, ZO1, and adherens junction markers, E-Cadherin and β-Catenin. While tight junctions are not affected, adherens junction proteins display interrupted cell boundary localization, with increased cytoplasmic localization. Scale bar: 5 µm. Ccdc66, coiled-coil domain-containing protein 66; IMCD3, inner medullary collecting duct cell line; PFA, paraformaldehyde; Arl13b, ADP-ribosylation factor-like protein 13B; DAPI, 4′,6-diamidino-2-phenylindole; ZO1, Tight junction protein ZO-1.

We defined spheroids (cell clusters of no more than 25 and less than 8 cells that formed a lumen in cross-section) as defective if they exhibited any of the following characteristics: failure to form a lumen, formation of multiple or distorted lumens in 3D, nuclei protruding into the lumen, disorganized distribution of apical and basal markers, or misoriented cilia (Fig. 7A). Analogous to the ciliogenesis defects of 2D serum-starved cultures, cilium length was reduced in Ccdc66-depleted cells grown in 3D cultures while ciliation efficiency was not affected (Fig. 7A, 7C). As assessed by fragmented DNA, we also found that CCDC66-depleted spheroids exhibited higher lumenal death than control spheroids (Fig. 7D).

Cell-to-cell adhesion is a critical process required for proper spheroid formation. Since we observed disturbed organization of apical, lateral and basal markers in 3D cultures, particularly the mislocalization of β-catenin, we further investigated cell-to-cell adhesion formation in 2D IMCD3 cultures. When grown to 100% confluency, IMCD3 cells typically form well-defined cell contacts with apico-basal polarity (Fig. 7E). However, CCDC66-depleted cells exhibited gross defects in cell-to-cell adhesion, as assessed by staining for E-cadherin and β-catenin, components of adherens junctions (Fig. 7E). These defects were less pronounced with the tight-junction marker ZO-1 (Fig. 7E). These results indicate that Ccdc66 plays an important role in maintaining cellular organization and adhesion in both 2D and 3D cultures, potentially due to its involvement in cilium maintenance, ciliary signaling and other non-ciliary functions.

## Discussion

CCDC66 is a MAP crucial for the proper function of microtubule-based structures, such as the cilium axoneme and the mitotic spindle, depending on the cell cycle stage. Its loss of function is directly linked to ciliopathies affecting the eyes and brain. Therefore, the previous research focused on the protein function in retinal pigment epithelium cells. Our findings highlight the significant role of Ccdc66 as a key regulator of cilium homeostasis and function in mouse kidney epithelial cells. We identified two specific roles for Ccdc66 at the primary cilium: First, it is required for maintaining the length, stability, and composition of steady-state and disassembling cilia. This involves regulating the stability of axonemal microtubules and cooperating with AurkA and HDAC6, two key regulators of cilium disassembly. Second, CCDC66 plays a role, either directly or indirectly, in activating the Hedgehog and Wnt signaling pathways, in part by regulating intraciliary transport. Importantly, both ciliary and potentially non-ciliary functions of Ccdc66 are necessary for tissue organization and cell polarity in 2D and 3D cultures. Our study provides important insights into the maintenance and disassembly of the ciliary axoneme and suggests that defects in cilium homeostasis, signaling, and tissue architecture could explain the pathogenesis of ciliopathies associated with CCDC66.

Our previous work showed that CCDC66 is required for efficiently assembling the primary cilium with the correct length and composition by cooperating with ciliary MAPs and transport complexes in human retinal epithelial cells [41, 42]. However, its roles in the maintenance and disassembly of the primary cilium, as well as in tissue organization, were unknown. We identified Ccdc66 as a critical regulator of these mechanisms using IMCD3 cells, which are suitable for investigating these processes. Stable expression of CCDC66 fluorescent function proteins in RPE1 cells showed localization to the ciliary tip and axoneme [42, 43]. However, this data could not be confirmed by endogenous staining due to the absence of specific antibodies. Our results provide the first confirmation of Ccdc66’s endogenous axoneme localization in IMCD3 cells. The loss-of-function studies in these cells reveal the role of Ccdc66 in cilium maintenance not assembly. The live cell imaging of ciliary dynamics further highlighted the role of CCDC66 in modulating the kinetics of cilium maintenance and disassembly. Specifically, it showed that CCDC66 depletion leads to fluctuations in ciliary length and increased ectocytosis during these phases. Moreover, its depletion leads to enhanced cilium disassembly, which was restored by inhibiting the AurkA-HDAC6 pathway that destabilizes axonemal microtubules. Taken together, we propose that CCDC66 acts as a microtubule-stabilizing factor, which regulates cilium maintenance and disassembly by stabilizing ciliary microtubules. Notably, CCDC66-expressing cilia were more resistant to axoneme destabilization by nocodazole treatment. We also showed that CCDC66 promotes formation of microtubule bundles that are resistant to dilution-induced disassembly *in vitro*. This mechanism nicely explains the frequent fluctuations in length of steady-state and disassembling cilia as well as enhanced cilium disassembly in Ccdc66-depleted cells. Given that cilium length is maintained by a balance between assembly and disassembly, the shorter cilium phenotype observed in IMCD3 cells could be explained by enhanced disassembly due to increased instability of axonemal microtubules without any defects in assembly.

MAPs stabilize microtubules through various mechanisms [4]. They do this by shifting the equilibrium toward polymerized tubulin, which involves slowing or inhibiting the dynamics of polymerized microtubules. They also promote stabilization by binding along the microtubule lattice, capping the ends to prevent depolymerization, and crosslinking microtubules with each other to create a more robust structure. CSPP1, a component of the ciliary tip module, was shown to inhibit microtubule growth and shortening, as well as stabilize damaged microtubule lattices [77]. A recent study showed that CCDC66, along with CEP104, CSPP1, TOGAGRAM1 and ARMC9, forms a complex at the ciliary tip which stabilizes microtubules by promoting slow and processive growth of the microtubule plus ends [50]. This study also found that CCDC66 does not affect the frequencies of catastrophes and rescues, nor did it induce pausing, suggesting that CCDC66 acts as a scaffold for the recruitment of the axonemal tip MAPs involved in regulating microtubule dynamics [50]. Notably, our findings that CEP104 cannot restore axonemal stability in CCDC66-depleted IMCD3 cells support this proposed mechanism. However, while the model nicely explains the behavior of singlet microtubule ends at the tip of cilia, it does not address the fact that these singlet microtubules are unstructured, intertwined, crosslinked bundles with actin filaments, adding complexity to ciliary tip structure [78, 79]. Given CCDC66’s direct affinity for actin and its microtubule bundling activity observed *in vitro*, as well as its localization not only to the ciliary tip but also along axonemal microtubules, we propose that it may stabilize microtubules by crosslinking microtubules with each other or with actin [40, 80]. Future research is necessary to test these models and elucidate how CCDC66 regulates microtubule stability *in* vitro.

Cilium disassembly involves a complex interplay among ciliary transport, cytoskeletal dynamics, and intracellular signaling [8, 81, 82]. Known regulators of cilium disassembly, including calcium, AurkA, HDAC6, and katanin, localize at the diverse cellular sites and function in various processes. This suggests that they might regulate cilium disassembly through multiple pathways. A similar multifaceted role may also apply to CCDC66. This is supported by the diverse interactions in cells revealed by the temporal proximity interactome of CCDC66 during cilium disassembly. While the CCDC66 disassembly proteome was enriched for proteins associated with the centrosome, cilium, and satellites, consistent with its cellular localization, other significant functional clusters related to cilium disassembly emerged, such as those involved in vesicular trafficking and lipid metabolism.

Considering cilium disassembly requires membrane remodeling in the ciliary pocket and the removal of ciliary receptors, CCDC66 may play a role in these processes [32, 82]. Additionally, actin-binding proteins were downregulated in the disassembly proteome compared to that of ciliated cells. Actin in cilia has been associated with ciliary ectocytosis and the myosin VI-dependent transport of proteins to the cilium [25, 65, 67]. Cortical actin also plays a critical role in regulating ciliary length [24]. These findings suggest that CCDC66 might regulate ectocytosis, actin-mediated trafficking or cortical actin by modulating actin dynamics during cilium disassembly and cilium length regulation. Future studies are needed to explore how CCDC66 influences cilium disassembly and length beyond its role in regulating microtubule stability.

Characterization of CCDC66 in both 2D and 3D IMCD3 culture models has revealed its roles in cell polarity and tissue organization, which depend largely on proper cilium homeostasis and function. Loss of CCDC66 results in shorter cilia and impairs ciliary signaling, suggesting that it may regulate polarity and tissue organization through its ciliary functions. Additionally, our results indicate that non-ciliary functions of CCDC66 could contribute to these processes. Disrupted cell polarity and apical cell-cell adhesion, specifically the adherens junctions (AJs), observed in both 2D and 3D cultures, suggest that CCDC66’s role in organizing the cytoplasmic microtubule and actin cytoskeleton could potentially underlie severe defects in epithelial organization. A similar role in apical junction formation has been identified for ciliopathy proteins within the NPHP8-NPHP4-NPHP1 module [83].

Depletion of these proteins led to irregular lumen formation in IMCD3 3D spheroids, while ciliation was largely unaffected, similar to our observation for CCDC66 [74, 83]. The significant defect in β-catenin distribution we observed in CCDC66-depleted cells may also suggest a role for CCDC66 in recruiting β-catenin to AJs. Since our analysis of genes downstream of β-catenin transcriptional activation suggests CCDC66’s role in the canonical Wnt signaling pathway, CCDC66 might also be involved in epithelial organization by affecting this pathway, in which β-catenin is the main effector. This is further supported by studies showing that impaired Wnt signaling contribute to cell death and defective cell polarity [84–86]. Future studies using 3D cultures of human colon carcinoma Caco-2 cells or MCF10A mammary epithelial cells, which do not form cilia, will be necessary to differentiate between the ciliary and non-ciliary contributions of CCDC66 to epithelial cell organization.

In conclusion, CCDC66 is emerging as a multifaceted regulator of ciliary and non-ciliary processes. *In vitro* and cellular studies have shown that CCDC66 regulates the abundance, organization and stability of microtubules in a context-dependent way [40–42, 80]. Our findings in this study define the ciliary functions of CCDC66 during maintenance and disassembly and suggest that both ciliary and non-ciliary roles contribute to epithelial cell organization. Together, these findings raise further questions about how CCDC66 regulates microtubules in its varied roles, which could either converge on microtubule stabilization or diversify through the regulation of different microtubule properties and interacting with other cellular players. A comprehensive understanding of CCDC66 mechanisms across its different cellular pools will advance our knowledge of its ciliary and non-ciliary roles.

Moreover, our study highlights the need to expand research on CCDC66 mechanisms beyond retinal cells and documented retinopathies to include potential abnormalities in kidney development and other epithelial tissue defects. This would enhance our understanding of CCDC66-linked ciliopathies and could influence the development of new treatments for these and related diseases.

## Materials and Methods

### Plasmids and Primers

Mouse Ccdc66 shRNA (sequence 5′-ATCAGGGATCCACTTCTTAAT-3′), targeting nucleotides 2635-2655 of mouse Ccdc66, was cloned into pLKO.1 (Invitrogen) at the EcoRI and AgeI sites. The control shRNA (sequence 5’-CAACAAGATGAAGAGCACCAA-3’) was previously described [87]. The plasmids pcDNA5.1-FRT/TO-miniTurboID-Flag-CCDC66 and pCDH-miniTurboID-Flag were previously described [42]. CEP104 cDNA (GenBank, NM_014704.4) was PCR amplified using forward primer 5’-GGGGACAACTTTGTACAAAAAAGTTGATatgccccacaagattggatttg-3’ and reverse primer 5’-GGGGACAACTTTGTACAAGAAAGTTGTgcgcttggcgtacgtcc-3’, cloned into pDONR221 and subsequently into pDEST-pcDNA5.1-FRT/TO-BirA*-Flag plasmid, which was provided by Anne-Claude Gingras (University of Toronto, Canada). Plasmids for protein expression in bacterial and insect cells, including pDEST-His-MBP, pDEST-His-MBP-mNeonGreen, pFastBac-HT-MBP-mNeonGreen-CCDC66, and pFastBac-HT-MBP-mCcdc66(clone 30626499), were previously described [40, 42, 88]. The pDONR221 plasmid containing the cDNA of human TOGARAM1 (GenBank, NM_001308120.2) was provided by Ronald Roepman (Radboud University Medical Center). The DNA fragment spanning the TOG1 and TOG2 domains of human TOGARAM1 (amino acids 2–621) was PCR amplified using primers forward 5’-GGGGACAACTTTGTACAAAAAAGTTGATctggcctcggccctc-3’, reverse 5’-GGGGACAACTTTGTACAAGAAAGTTGTtcaactctgagttctgttaccagcc-3’, and subsequently cloned into pDONR221 and then into pDEST-His-MBP vector for bacterial expression.

### Cell culture, transfection and transduction

Mouse Inner medullary collecting duct cell line (IMCD3) cells were cultured in Dulbecco’s modified Eagle’s Medium DMEM/F12 50/50 medium (Pan Biotech, Cat. # P04-41250) supplemented with 10% Fetal Bovine Serum (FBS, Life Technologies, Ref. # 10270-106, Lot # 42Q5283K) and 1% penicillin-streptomycin (GIBCO, Cat. # 1540-122). Human embryonic kidney (HEK293T, ATCC, CRL-3216) cells were cultured with DMEM medium (Pan Biotech, Cat. # P04-03590) supplemented with 10% FBS and 1% penicillin-streptomycin. All cell lines were authenticated by Multiplex Cell Line Authentication (MCA) and were tested for mycoplasma by MycoAlert Mycoplasma Detection Kit (Lonza). HEK293T cells were transfected with the plasmids using 1mg/ml polyethylenimine, MW 25 kDa (PEI).

Recombinant lentiviruses were generated in HEK293T cells using pLKO.1 Scramble shRNA, pLKO.1 msCCDC66 shRNA, pCDH-miniTurboID-Flag plasmids as transfer vectors co-transfected with packaging and envelope vectors, psPax2 (Addgene #12260) and pMD2.G (Addgene #12259) using 1 mg/ml polyethylenimine, MW 25 kDa (PEI). 48Dh after transfection, the supernatant was harvested, filtered, and titered on IMCD3 cells using GFP-expressing lentivirus. For transduction, IMCD3 cells were plated in 6-well tissue culture plates at 500 × 10^5^, infected with an approximate 1 MOI over 2 days. For short-term depletion with shRNAs and stable long-term expression of fusion proteins, cells were selected with a medium containing 3 μg/mL puromycin medium (Invitrogen, CA) for 4 days, then assayed 6–10 days after initial infection. The heterogeneous pools were used for functional assays. Depletion of the Ccdc66 mRNA and protein was confirmed by immunofluorescence, immunoblotting and qPCR.

### 3D spheroid assay

40 μl/well of 100% Matrigel (BD Biosciences) was solidified in Lab-Tek 8-well chamber slides (Thermo Fisher) by incubating for 15 min at 37°C. Control and Ccdc66-depleted IMCD3 cells were trypsinized, washed with PBS and resuspended in DMEM/F12 50/50 supplemented with 2% fetal bovine serum and 2% Matrigel.

Then, 5,000 cells/well were plated in Matrigel-coated 8-well chamber slides. Cells were cultured at 37°C for 4 days, serum-starved for 2 days, fixed with 4% PFA and processed for immunofluorescence by staining with the indicated markers.

### Protein expression and purification

Protein expression and purification were performed as previously described [40]. BEVS baculovirus expression system and protocol was used for the expression of tagged full-length CCDC66 protein [89]. Briefly, 100 mL of Hi5 cells (1 x 10^6^ cell/mL) were infected with P1 baculovirus produced in Sf9 cells, carrying His-MBP-mNeonGreen-CCDC66, at MOI of 1. Cells were collected 48 hr after infection. For protein purification, bacterial or insect cells were lysed by sonication in lysis buffer (20 mM Hepes, pH 7.0 (or pH 7.5 for full-length protein), 250 mM NaCl, 0.1% Tween20, 2 mg/ml lysozyme, 1 mM PMSF, 1mM protease inhibitor cocktail, 5mM BME and 10 mM Imidazole). His-tagged proteins were subsequently purified using Ni-NTA–agarose beads (Thermo Scientific) and elution buffer (20 mM HEPES, pH 7.0 or pH7.5, 250 mM NaCl, 5mM BME, and 250 mM imidazole). For subsequent microtubule assays, proteins were dialyzed against BRB80 buffer (80 mM PIPES pH 6.8, 1 mM EGTA, 1m M MgCl2). For subsequent antibody production, proteins were dialyzed against PBS.

### *In vitro* microtubule stabilization assay

Stability assay by dilution was performed as previously described snc.) at a 10:1 ratio in BRB80 with 1 mM DTT, 1mM GTP and 25% glycerol for 30 min at 37°C. Following glycerol polymerization, MT were either 10x diluted in BRB80 buffer with only DTT (no GTP and glycerol), or buffer with 100 nM of protein His-MBP-mNeonGreen, His-MBP-mNeonGreen-CCDC66 or His-MBP-TOGARAM1_TOG1TOG2_. After 5 min incubation, reaction mixtures were fixed with 1% glutaraldehyde in BRB80 with DTT for 3 min, quenched with 100 mM Tris pH8.8 (final concentration) for 1 min, totaling to 11x dilution, then only half of the reaction was pelleted onto poly-L-lysine coated glass coverslips. MT control before dilution was immediately fixed with 1% glutaraldehyde in the volume of BRB80 with DTT, GTP, and glycerol to make the same 11x dilution, and half of the reaction was pelleted together with diluted samples. Centrifugation of fixed MT reactions was performed at 4700 rpm for 45 min above 40% glycerol BRB80 cushion, fixed on coverslips with ice-cold methanol, and processed for immunofluorescence using anti-Tubulin antibody and confocal microscopy. The experiment was repeated 3 times.

### Ultra-Structure Expansion Microscopy (U-ExM)

U-ExM was performed as previously described [92] Briefly, IMCD3 cells were grown on glass coverslips and serum starved for 48 h. Coverslips were incubated in 1.4% formaldehyde / 2% acrylamide ( 2X FA / AA) solution in 1X PBS for 5 h at 37°C before gelation in Monomer Solution supplemented with TEMED and APS (final concentration of 0.5%) for 1 hr at 37°C. Denaturation was performed at 95°C for 1h 30 min and gels were stained with primary antibodies for 3 h at 37°C. Gels were washed three times for 10 min at RT with 1 X PBS with 0.1% Triton-X (PBS-T) before secondary antibody incubation for 2h 30 min at 37°C followed by three 10 min washes in PBS-T at RT. Gels were expanded by three exchanges of 150 ml dH_2_O before imaging. The following reagents were used in U-ExM experiment: formaldehyde (FA, 36.5–38%, F8775, Sigma-Aldrich), acrylamide (AA, 40%, A4058, Sigma-Aldrich), N,N’-methylenbisacrylamide (BIS, 2%, M1533, SIGMA), sodium acrylate (SA, 97–99%, 408220, Sigma-Aldrich), ammonium persulfate (APS, 17874, Thermo Scientific), tetramethylethylendiamine (TEMED, 17919, Thermo Scientific), and poly-D-Lysine (A3890401, Gibco).

### Immunofluorescence, antibodies and drug treatments

Cells were grown on coverslips, washed with PBS, and fixed by either 4% PFA diluted in cytoskeleton buffer (100 mM NaCl (Sigma-Aldrich, S9888), 300 mM sucrose (Sigma-Aldrich, S0389), 3 mM MgCl2 (Sigma-Aldrich, M2670), and 10 mM PIPES (Sigma-Aldrich, P6757)) for 10 min at room temperature or ice-cold methanol at −20°C for 10 minutes. After washing three times with PBS, cells were blocked with 3% BSA (Capricorn Scientific) in PBS plus 0.1% Triton X-100 and incubated with primary antibodies in blocking solution for 1 h at room temperature. Cells were washed three times with PBS and incubated with secondary antibodies at 1:2000 dilution for 1 h and DAPI (Thermo Fisher Scientific cat#D1306) at 1:1000 for 5 min at room temperature. Following three washes with PBS, cells were mounted using Mowiol mounting medium containing N-propyl gallate (Sigma-Aldrich). Anti-CCDC66 antibody was generated by immunizing rats (Koc University, Animal Facility) with His-MBP-tagged mouse CCDC66 (clone 30626499) comprising amino acids 1–756 purified from Hi5 insect cells. The antibody was affinity purified against His–MBP– mCcdc66 (a.a. 1–756) and used at 0.5 μg/ml for immunofluorescence. Other primary antibodies used for immunofluorescence were mouse anti-acetylated tubulin (clone 6-11B, 32270, Thermo Fisher) at 1:10000, mouse anti-polyglutamylated tubulin (AG-20B-0020, clone GT335, Adipogen) at 1:1000; mouse anti gamma-tubulin (Sigma, clone GTU-88, T5326) at 1:1000, mouse anti-alpha-tubulin (Sigma-Aldrich, DM1A) at 1:1,000, rabbit anti-ARL13B (17711-1-AP, Proteintech) at 1:250, mouse anti-ARL13B (75-287, NIH Neuromab) at 1:250, rabbit anti-Flag (CST #2368) at 1:200, rabbit anti-CEP164 (Proteintech, 22227-1-AP) at 1:200, rabbit anti-KIAA0568/Talpid3 (Proteintech 24421-1-AP) at 1:100, rabbit anti-CP110 (A301-344A, Betyl) at 1:500, goat anti-PCM1 (sc-50164, Santa Cruz Biotechnology), anti-Smoothened (sc-166685, Santa Cruz Biotechnology) at 1:50, rabbit anti-IFT88 (Proteintech 13967-1-AP) at 1:100, rabbit anti-BBS4 (Proteintech 12766-1-AP) at 1:100, mouse anti-E-cadherin (Proteintech 60335-1-Ig) at 1:100, rabbit anti-β-Catenin (ProteinTech PTG 51067-2-AP) at 1:200, mouse anti-ZO1 (Invitrogen 33-9100) at 1:1000, Phalloidin-AlexaFluor 568 coupled at 1:1500, Streptavidin-AlexaFluor 568 coupled at 1:2000.

Secondary antibodies used for immunofluorescence experiments were AlexaFluor 488-, 568- or 633-coupled (Life Technologies) and they were used at 1:2000. For microtubule depolymerization experiments, cells were treated with 400 ng/ml nocodazole (Sigma-Aldrich, Cat. #M1404) or vehicle (dimethyl sulfoxide) for 2 h at 37°C. AurkA and HDCA6 inhibition was performed with 500 nM MLN8237 or 2 µM Tubacin for 6 h at 37°C. Smoothened activation was induced by 200 nM SAG (EMD Millipore) treatment for 4 h in serum-depleted IMCD3 cultures.

### Microscopy and Image analysis

For the assessment of protein localization and level quantifications, images were acquired with Leica DMi8 fluorescent microscope with a stack size of 5 µm and step size of 0.25 µm in 1024×1024 format using an HC PL APO CS2 63×1.4 NA oil objective. Higher resolution images were taken by using an HC PL APO CS2 63×1.4 NA oil objective with Leica SP8 confocal microscope 1024×1024 pixel format, with pixel size ranging from 90 to 45 nm. For imaging spheroids, images were acquired with HC PL APO CS2 63×1.4 NA oil objective with Leica SP8 confocal microscope with a stack size of 50-70 µm and step size of 0.5 µm in 1024×1024 format with pixel size of 150 nm (above Nyquist sampling rate). For super-resolution imaging of endogenous CCDC66 localization, images were acquired using Elyra 7 with Lattice SIM² (Zeiss) and Zeiss Objective Plan-Apochromat 63×/1.4 Oil DIC M27, 633 nm, 568 nm and 488 nm laser illumination, and standard excitation and emission filter sets. Sections were acquired at 0.110 μm z-steps. U-ExM samples were imaged using a Leica STELLARIS8 confocal microscope equipped with an HC PL APO CS2 63×1.4 NA oil objective, and the images were denoised using the Lightning Wizard. Time-lapse live imaging was performed with Leica SP8 confocal microscope equipped with an incubation chamber using HC PL APO CS2 63x 1.4 NA oil objective. To image the effect of CCDC66 depletion on cilium maintenance, control or Ccdc66-depleted IMCD3::GFP-SSTR3 cells were seeded at 1×10^5^ cells/ml to FluoroDishes (WPI Europe). The following day, they were serum starved in DMEM/F12 without FBS for 48 h and subsequently imaged at 37°C with 5% CO2. Imaging was performed with a frequency of 4 minutes per frame, with a 1 μm step size and a 12 μm stack size, in a 512×512 pixel format for 6 h. For observing cilium disassembly, control and depleted cells were serum-starved for 48 h. After serum stimulation, cilium disassembly was imaged every 4 minutes for 6 h in a 512×512 pixel format. Images were processed using ImageJ (National Institutes of Health, Bethesda, MD).

The percentage of ciliated cells was determined by counting the total number of cells and the number of cells with primary cilia, as identified through DAPI staining of nuclei and either Arl13b or acetylated tubulin immunofluorescence of primary cilium. For quantitative immunofluorescence of ciliary and centrosomal protein levels, z-stacks of cells were acquired using identical gain and exposure settings, which were determined based on the fluorescence signal in control cells. These z-stacks were then used to assemble maximum-intensity projections using ImageJ (National Institutes of Health, Bethesda, MD). Ciliary regions were identified in these images by the presence of Arl13b and/or acetylated tubulin signals. The centrosome regions in these images were defined by centrosomal marker staining for each cell, and the total pixel intensity of a circular 1.5 μm^2^ area centered on the centrosome in each cell was measured using ImageJ and defined as the centrosomal intensity. Background intensity was subtracted from the centrosomal and ciliary fluorescence intensities. This subtraction was performed by quantifying fluorescence intensity in a region of equal dimensions in the area adjacent to the primary cilium or centrosome. Ciliary protein concentration was determined by dividing the fluorescence signal of the protein to the cilium area, which was quantified using Arl13B or acetylated tubulin staining. Ciliary protein concentrations and centrosomal protein levels were normalized relative to the control group’s mean (=1). Statistical analysis was done by normalizing these values to their mean. Quantification of Cp110 localization at the centrosome was performed by counting the total number of centrosomes, identified using a gamma-tubulin marker, and the number of centrosomes exhibiting Cp110 signal. The percentage of centrosomes positive for Cp110 was then calculated.

Spheroids were manually analyzed to determine lumen formation and cell polarity based on staining for F-actin and β-catenin staining. Ciliary defects were assessed using Arl13b staining. Cell clusters smaller than 8 cells in cross-section or larger than 50 cells without a lumen were excluded from the analysis. With these criteria, approximately 50% of IMCD3 cells successfully formed spheroids. Among these, defective spheroids were defined as the ones that lacked a hollow lumen, had multiple or distorted lumens, showed disorganized apical and basal markers, and/or exhibited misaligned nuclei protruding into the lumen using the 3D rendering tool of LasX software (Leica Microsystems). Due to a large step size of 0.5 µm, which captured the entire size of the spheroids but omitted some cilia length details, cilia length measurements were not possible. Cilia count was based on Arl13b staining within regions stained for F-actin and β-catenin and was normalized by total cell count determined by DAPI staining to calculate the frequency of luminal cilia. Cell death was identified by counting apoptotic bodies (fragmented, condensed DNA based on DAPI intensity) and was limited to those within the spheroid lumen. This count was divided by the total cell number to calculate the frequency of luminal cell death.

For the analysis of live imaging movies, ciliary length was measured using a freehand line tool on 3D-rendered images in ImageJ. The quantification of ectocytosis events was performed manually by analyzing each cilium. The tips and bases of the cilia were identified using the cell’s cytoplasm, marked by GFP-SSTR3, as a reference for the base. Fragments and vesicles were lost during imaging, while the cilia remained associated with the cell through its base. Fragment ectocytosis was differentiated from vesicle ectocytosis by the size of the released ciliary piece; vesicle ectocytosis involved vesicles smaller than 0.5 μm, while fragment ectocytosis involved fragments larger than 0.5 μm.

### Cell lysis and immunoblotting

Cells were lysed in RIPA buffer and protease inhibitors for 30 min at 4°C followed by centrifugation at 10,000 rpm for 30 min. The protein concentration of the resulting supernatants was determined with the Bradford solution (Bio-Rad Laboratories, CA, USA). For immunoblotting, equal quantities of cell extracts were resolved on SDS-PAGE gels, transferred onto nitrocellulose membranes and blocked with TBS with 0.1% Triton-X100 with 5% milk for 1 h at room temperature. Blots were incubated with primary antibodies diluted in 5% BSA in TBS-TX overnight at 4°C, washed with TBS-TX three times for 10 minutes and blotted with secondary antibodies for 1 h at room temperature. After washing blots with TBS-TX three times for 5 minutes, they were visualized with the LI-COR Odyssey® Infrared Imaging System and software at 169 mm (LI-COR Biosciences). Primary antibodies used for immunoblotting were rat anti-Ccdc66(homemade) at 1:50, mouse anti-Gapdh (CST #97166) at 1:1000, mouse anti-acetylated tubulin (clone 6-11B, 32270, Thermo Fisher) at 1:10000, rabbit anti-GFP at 1:2000 (homemade, previously described in [63]), anti-Flag (CST #2368) at 1:1000, Streptavidin-IRDye800 coupled at 1:10000. Secondary antibodies used for western blotting experiments were IRDye680- and IRDye800-coupled and were used at 1:15000 (LI-COR Biosciences).

### RNA isolation, cDNA synthesis and qPCR

Total RNA was collected from control and CCDC66-depleted cells serum starved for 48 h to induce cilia formation. For Hedgehog pathway analysis, cells were also treated with 200 nM SAG 24 h before collection. RNA was isolated using the NucleoSpin RNA kit (Macherey-Nagel) according to the manufacturer’s protocol. The quantity and purity of RNA were assessed by measuring the optical density at 260 and 280 nm. Single-strand cDNA synthesis was performed with 1 µg of total RNA using SCRIPT Reverse Transcriptase (Jena Bioscience Cat#: PCR-511S) or iScript cDNA synthesis kit (BioRad, # 1708891). qPCR analysis of Ccdc66 mRNA levels was performed using GoTaq® qPCR Master Mix (Promega) with primers 5′-GCAGAAAGCTGCCACAGAGA-3′ and 5′-CTGGGCTCTTCTTGCTTCCA-3′.

Mouse Gapdh or β-actin was used as normalization controls with primers 5’-AAGGTCATCCCAGAGCTGAA – 3’ and 5’ – CTGCTTCACCACCTTCTTGA – 3’ or 5’ – GTTCGCCTTCATTATGGACTGCC-3’ and 5’ – ATAGCACCCTGTTCCCGCAAAG-3’, respectively. Components of the Hedgehog signaling pathway, Gli1 and Ptch1, were analyzed using primers 5′-GCATGGGAACAGAAGGACTTTC-3′ and 5′-CCTGGGACCCTGACATAAAGTT-3′ and primers 5′-TGAACTGGGCAGCTATGAAGTC-3′ and 5′-ATGCTCCTTTCCTCCTGAAACC-3′, respectively. Downstream effectors of Wnt signaling using primers 5’ – TGCGTTCTCGGAATAGCTCC-3’ and 5’ – AGAGCTTTGCTGTAAAAGAGAGGA – 3’ for Axin2, 5’ – GGGCCAGGTGGACTTCTATG – 3’ and 5’ – CCACCGAACAATAAGGAGTGC-3’ for Pdk2 and 5’ – AACTGGTTACCCTTCCACACC – 3’ and 5’ – TCCAGGAACTTGGAACTGTAAGG-3’ for Fibronectin.

### Biotin identification experiments and mass spectrometry data analysis

IMCD3 cells stably expressing Flag-miniTurbo or Flag-miniTurbo-CCDC66 previously described [42] were used. For mass spectrometry analysis, each cell type was grown in 5×15 cm plates in DMEM/F12 medium supplied with 10% FBS and 1% penicillin-streptomycin. The ciliated cell populations were generated after growing cells to 100% confluency and serum starving them for 48 h in DMEM/F12 with 0% FBS, followed by incubation with 500 μM biotin for 30 min. Cilium disassembly was induced in cells that had been serum-starved for 48 h, followed by serum stimulation for either 1 or 6 h. For biotinylation, cells undergoing disassembly were treated with 500 μM biotin for the last 30 minutes of the serum stimulation period. After the biotin treatment, cells were washed twice with PBS and lysed using RIPA buffer (50 mM Tris-HCl pH 8.0, 150 mM NaCl, 0.1% SDS, 0.5% sodium deoxycholate, 1% Triton X-100), which was freshly supplemented with a protease inhibitor cocktail and 1 mM PMSF. Cell lysates were sonicated and then centrifuged at 16,000 g for 1 hour at 4°C. The resulting supernatant was incubated with Streptavidin–agarose beads (Thermo Fisher Scientific) for 16 h at 4°C. After incubation, beads were washed twice with the RIPA buffer, once with 1 M KCl, once with 0.1 M Na_2_CO_3_, once with 2 M urea in 10 mM Tris-HCl pH 8.0 and finally twice with the RIPA buffer. For mass spectrometry analysis, the beads were resuspended in 100 µl of 50 mM ammonium bicarbonate and and analyzed at the KUPAM proteomics facility as previously described [87]. The data presented in Figure 5, Figure S4, and Table S1 were obtained from two biological and two technical replicates each for Flag-miniTurbo CCDC66 ciliation, Flag-miniTurbo ciliation, and Flag-miniTurbo 1-hour disassembly.

For the Flag-miniTurbo CCDC66 1-hour and 6-hour disassembly, data were derived from two biological replicates, one with two technical replicates and the other with three technical replicates.

For mass spectrometry analysis, Normalized Spectral Abundance Factor (NSAF) values were calculated for each protein by dividing each Peptide Spectrum Match (PSM) by the total PSM count in that dataset. The datasets were filtered using several steps. First, proteins present only in the control dataset and in just one of the technical replicate datasets were excluded. NSAF values in the CCDC66 datasets were divided by the corresponding NSAF values from the control dataset to calculate an enrichment score, referred to as fold change hereafter. The average fold change was calculated for proteins present in both biological replicates. Proteins with an average log_2_(Fold Change) score less than 1 were removed. Next, the remaining proteins were analyzed using CRAPome (https://reprint-apms.org), a contaminant repository for mass spectrometry data from affinity purification experiments, where a list of contaminancy percentages (%) were generated [93]. Proteins with a contamination percentage greater than 50% were considered contaminants and removed. This cutoff value was selected based on the presence of any known interaction partners of CCDC66 within that range. Finally, proteins were organized according to their log_2_(Fold Change) values and an interactome map was generated using Cytoscape [94]. The edges of the high confidence hit proteins in the proximity map were plotted with the STRING database, and the map was visualized by CytoScape [95]. The functional clusters and GO categories for these clusters were determined with the Database for Annotation, Visualization and Integrated Discoverey (DAVID) with a significance threshold of P < 0.05, and supplemented by literature mining and literature mining [96].

### Quantification and statistical analysis

Data were analyzed and plotted using GrapPad Prism 9 (GraphPad, La Jolla, CA). Results were presented as mean ± standard error of the mean (SEM) or mean ± standard deviation (SD). The number of biological replicates was indicated in the figure legends. Two-tailed unpaired t-tests, unpaired t-tests with Welch correction (Welch’s t-test), one-sample t-tests, and one-way analysis of variance (ANOVA) were used to assess the statistical significance of measurements with a normal distribution. Data that did not follow a normal distribution were analyzed using a non-parametric Wilcoxon test. Normality was assessed using the Shapiro-Wilk Test. The following key is used for asterisks indicating p-values in the figures: *p < 0.05, **p < 0.01, ***p < 0.001, ns indicates not significant.

## Acknowledgements

We acknowledge Melis Dilara Arslanhan, Ezgi Odabasi and Efe Begar for their insightful feedback on this work. We thank Melis Dilara Arslanhan Gül for help with UxEM and Serra Doganata for help with data analysis. We also acknowledge use of the services and facilities of the KUPAM-Koç University Proteomics Facility. This project has received funding from the European Union’s Horizon 2020 research and innovation program under the Marie Sklodowska-Curie grant agreement No 896644 awarded to JD and European Union’s Horizon 2020 and Europe research and innovation program under the European Research Council Starting grant agreement CentSatRegFunc-679140 and SatellieteHomeostatis - 101078097 to ENF. This work was also supported by EMBO Installation Grant 3622 and Young Investigator Award to ENF, TUBITAK BIDEB 120C148 grant to ENF and TUBITAK ARDEB to 120Z179 to JD. Fig. 5C was created with BioRender.com.

## Competing interest

The authors declare no competing interests.

---

**Table 1.** Mass spectrometry results related to Figure 5 and Figure S4. Column explanations were placed to sheet named “Legend”.

## Supplementary Figures

**Figure S1.**
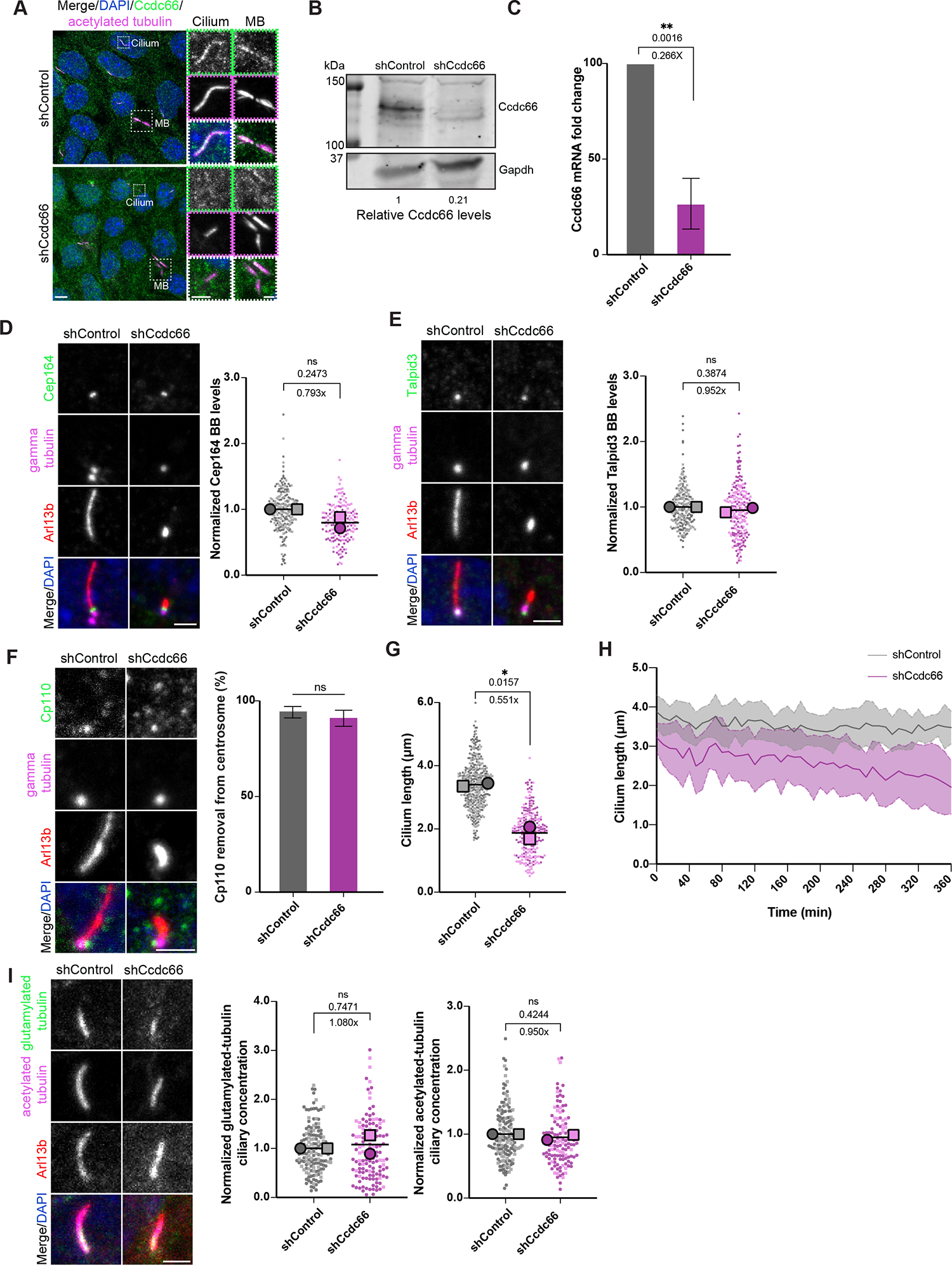
Ccdc66 regulates cilium stability independently of the basal body ciliogenesis factors and axonemal PTMs. (A-C) Ccdc66 was successfully depleted from IMCD3 cells. Control and Ccdc66-depleted cells were serum starved for 48h, fixed with 4% PFA and stained against CCDC66 with homemade antibody, anti-acetylated-tubulin, and DAPI. Scale bars: 5 µm. Insets show 3× magnifications of the cilia and midbody signals. Scale bar 2 µm (B) Western blot analysis of cell lysates from control and Ccdc66-depleted samples using homemade Ccdc66 antibody and mouse anti-Gapdh as loading control. Represented is the fold change of band intensities normalized to loading control of displayed experimental replicates. n=2 (C) qPCR analysis of mRNA isolated from control and Ccdc66-depleted cells with primers recognizing C-terminal internal region and Gapdh primers as normalization control. Box plot shows mean ± SD. n=4 (One sample t test p value **p=0.0016) (D-F). Control and Ccdc66-depleted cells were ciliated for 48h, fixed with methanol and stained against anti-CEP164, anti-gamma tubulin to mark basal bodies, and anti-Arl13b. The super plot of normalized individual experimental values with mean± SEM represents 2 independent experiments. The experimental replicates, represented by circles or squares, display varying shades to differentiate between different replicates. n > 150 cells for each condition. The mean basal body fluorescence intensity of Cep164 in Ccdc66 depletion is ∼0.8-fold decreased compared to the control mean. (Welch’s t test p value ns: not significant) (E) Fluorescence intensity levels of Talpid3 at the basal body were assed as in (D). n > 200 cells for each condition. The mean basal body fluorescence intensity of Talpid3 in Ccdc66 depletion is 0.95-fold of the control mean. (Welch’s t test p value ns: not significant) (F) The removal of Cp110 from centrosomes, from both ciliated and non-ciliated cells, was quantified using gamma-tubulin as the centrosome marker. n> 200 cells for both conditions. (G) Raw data of cilia length quantification in Figure 1D. (H) Raw data of cilia length quantification in Figure 1E. (I) Quantification of ciliary axoneme PTMs in control and Ccdc66-depleted cells, serum starved and stained against acetylated and polyglutamylated-tubulin and Ar13b to mark cilia. Measured are ciliary integrated densities=mean intensity*area, to better represent levels of PTMs in cilia of varying size. The super plot represents normalized individual experimental values and means ± SD of 2 independent experiments. n > 150 cilia for each conditioned and PTM. (p value of Welch’s t tests ns: not significant) Ccdc66, coiled-coil domain-containing protein 66; IMCD3, inner medullary collecting duct cell line; PFA, paraformaldehyde; DAPI, 4′,6-diamidino-2-phenylindole; qPCR, quantitative polymerase chain reaction; mRNA, messenger ribonucleic acid; Gapdh, Glyceraldehyde 3-phosphate dehydrogenase; Arl13b, ADP-ribosylation factor-like protein 13B; Cep164, Centrosomal protein of 164 kDa; CP110, Centriolar coiled-coil protein of 110 kDa; PTM, posttranslational modification.

**Figure S2.**
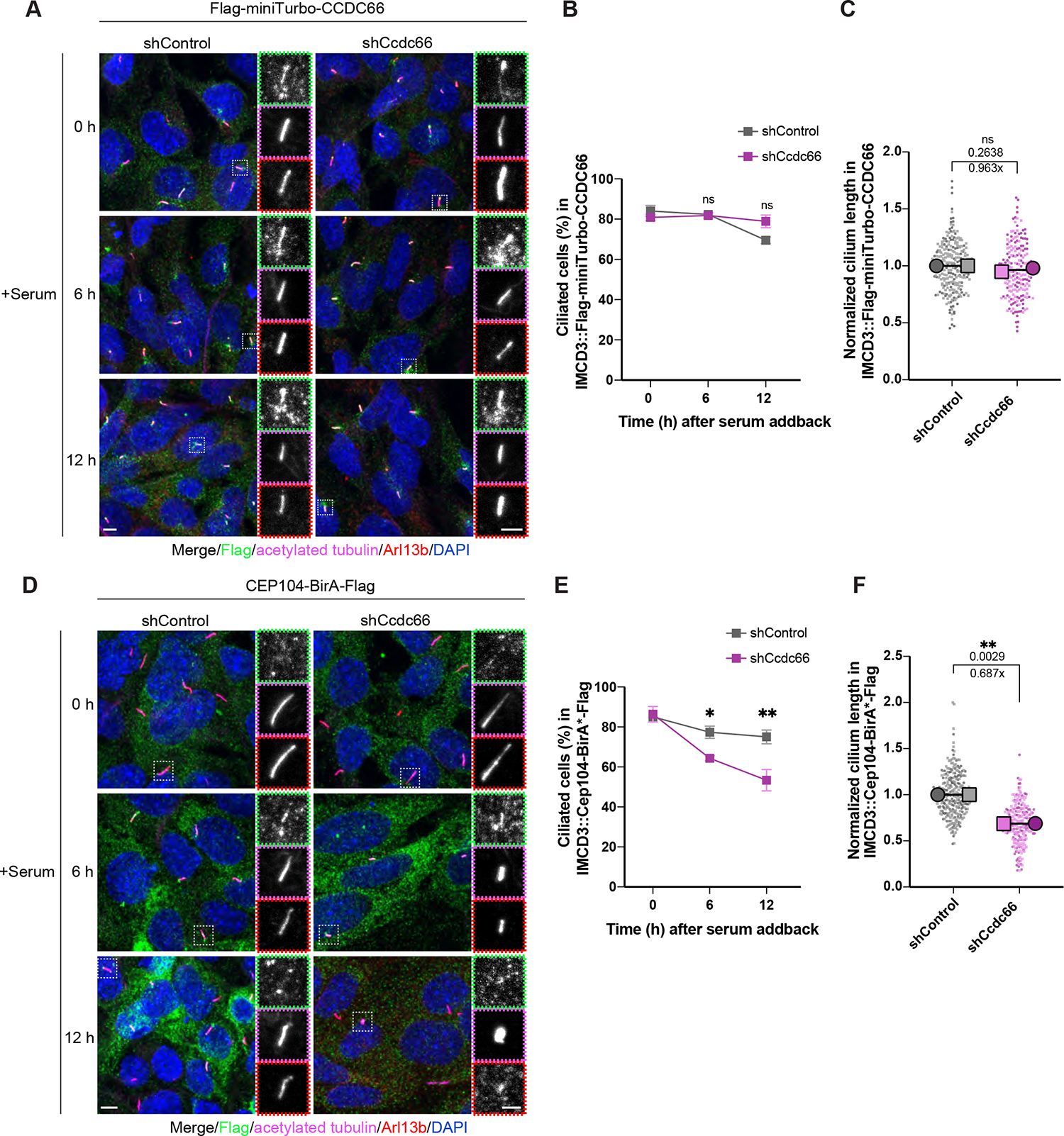
Cilium disassembly defects were rescued by ectopic expression of Ccdc66, but not by Cep104. (A-C) IMCD3::Flag-miniTurbo-CCDC66 cells were transduced with control and Ccdc66-targeting shRNA, selected and seeded on coverslips. After 48 h serum starvation, the cells were stimulated with serum-rich medium to induce cilia disassembly for total of 12 h, fixed and stained with anti-Flag, anti-acetylated-tubulin, anti-Arl13b and DAPI. Scale bar: 5 µm. Insets show 3× magnifications of the cilia, Scale bar: 2 µm (B) Quantification of cilia number in (A). Data represents mean ± SD of 2 independent experiments. n > 400 cells for control and CCDC66 depletion at all three indicated time points. The mean cilia percentage at 0 h is 84.01 for shControl, 80.82% for shCCDC66; at 6 h is 82.31% for shControl, 81.84% for shCCDC66; at 12 h is 69.48% for shControl, 78.90% for shCcdc66. (p values of multiple t test of grouped data p=0.2853, p=0.7710, p=0.0645). (C) Quantification of cilia length in (A) at 0 h. Super plot represents normalized individual experimental values with mean ± SEM of 2 independent. n > 400 cells for each condition. The mean cilia length in Ccdc66 depletion is 0.96-fold decreased compared to the mean control length. (Welch t test p value ns: not significant). (D) IMCD3::CEP104-BirA*-Flag were transduced with control and Ccdc66-targeting shRNA, selected and seeded on coverslips. After 48 h serum starvation, followed by serum addback for total of 12 h, fixed and stained with anti-Flag, anti-acetylated-tubulin, anti-Arl13b and DAPI. Scale bar: 5 µm. Insets show 3× magnifications of the cilia, Scale bar: 2 µm (E) Quantification of cilia number in (D). Data represents mean ± SD of 2 independent experiments. n > 400 cells for control and CCDC66 depletion at all three indicated time points. Mean cilia percentage at 0 h is 85.09% for shControl, 86.38% for shCCDC66; at 6 h is 77.37% for shControl, 64.34% for shCcdc66; at 12 h is 75.06% for shControl, 53.36% for shCcdc66. (p values of multiple t test of grouped data p=0.6415, **p=0.0085, **p=0.0066) (F) Quantification of cilia length in (D) at 0 h. The super plot represents normalized individual experimental values with mean ± SEM of 2 independent. n > 400 cells for each condition. The mean cilia length in CCDC66 depletion is 0.69-fold decreased compared to the mean control length. (Welch’s t test p value **p=0.0029) Ccdc66, coiled-coil domain-containing protein 66; IMCD3, inner medullary collecting duct cell line; shRNA, short hairpin RNA; DAPI, 4′,6-diamidino-2-phenylindole; Arl13b, ADP-ribosylation factor-like protein 13B; CEP104, Centrosomal protein of 104 kDa; SEM, standard error of mean; SD, standard deviation.

**Figure S3.**
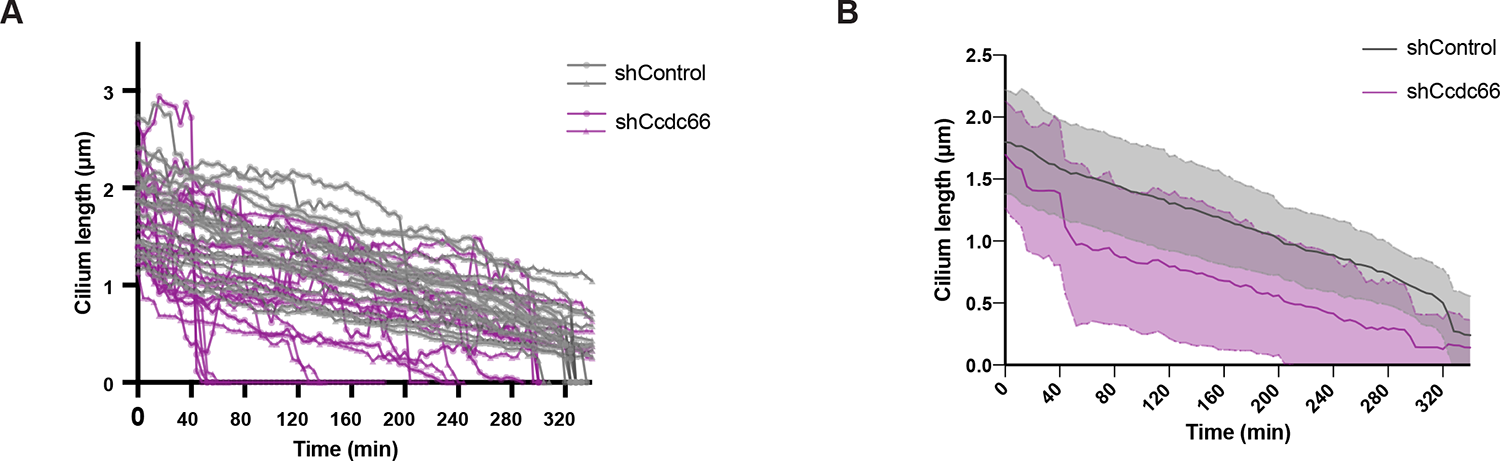
Live-cell imaging demonstrates that cilia disassembly predominantly occurs through combined and instant cilia loss in Ccdc66-depleted cells. (A, B) Raw cilia length curves from Figure 3B, measured from fluorescence of two independent experiments and represented as the mean ± SD. n= 20 cilia for both shControl and shCcdc66 conditions. CCDC66, coiled-coil domain-containing protein 66; IMCD3, inner medullary collecting duct cell line; SSTR3, Somatostatin receptor type 3; GFP, green fluorescent protein; SEM, standard error of the mean; SD, standard deviation.

**Figure S4.**
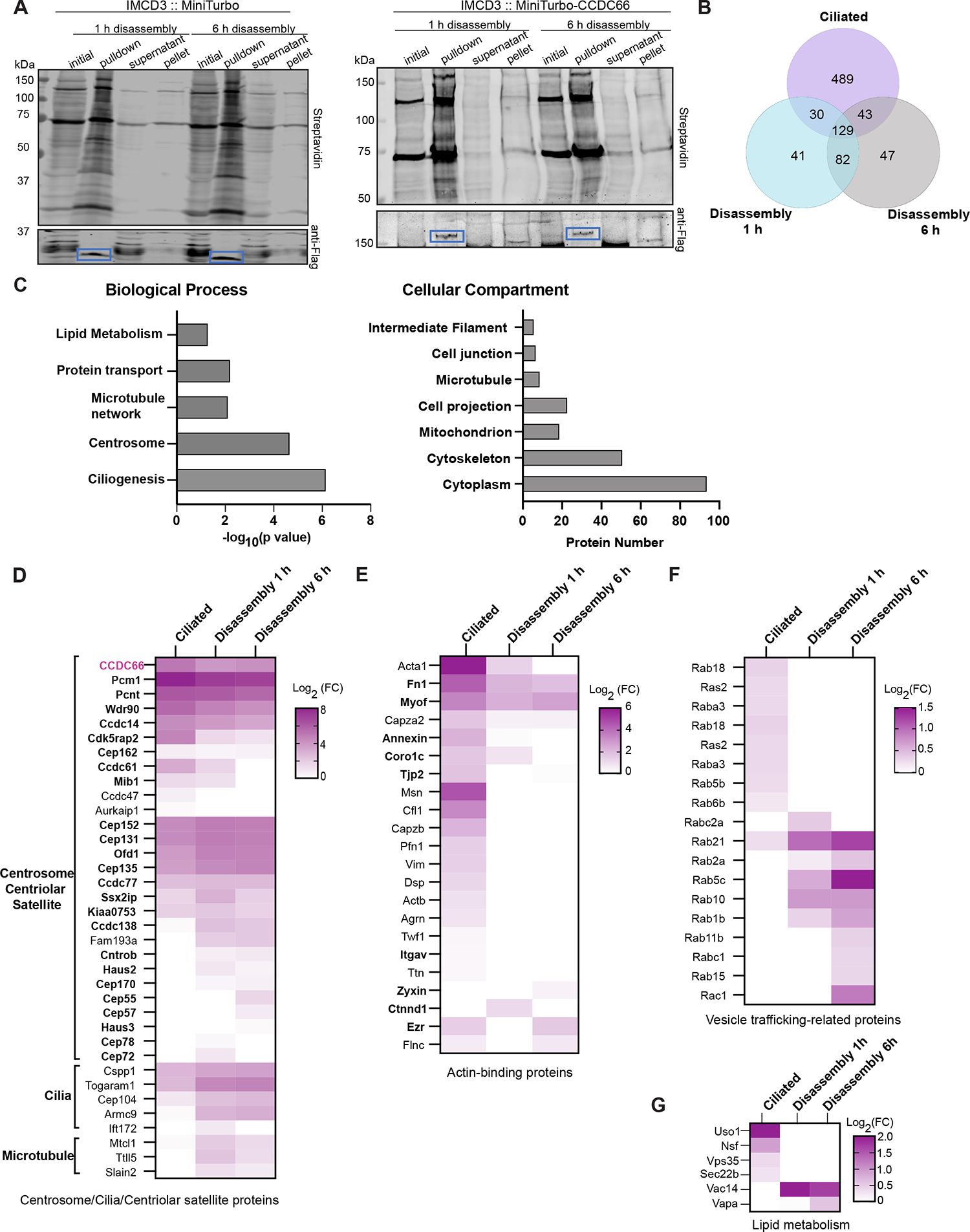
GO and comparative analysis of CCDC66 ciliated and disassembly proximity interactomes (A) Western blot analysis of samples from different stages of streptavidin pulldowns from IMCD3 cells expressing Flag-miniTurboID or Flag-miniTurboID-CCDC66. Cells were ciliated by serum starvation and then stimulated with serum for 1 h and 6 h. Samples were collected at the lysate (initial), pellet, and supernatant stages, and the non-eluted bead samples were run as the pulldown samples. Samples were blotted using Streptavidin-IRDye800 coupled and anti-Flag. Blue boxes show bands corresponding to Flag fusion proteins. (B) Comparison of the Ccdc66 proximity interactomes of ciliation, 1 h disassembly and 6 h disassembly conditions. (C) GO-enrichment analysis of the Ccdc66 disassembly interactome based on their biological process and cellular compartment. The x-axis represents the log-transformed p-value (Fisher’s exact test) of GO terms. (D-F) Heat map showing Log_2_(Fold Change) of the Ccdc66 proximity interactors in ciliated cells versus 1 hour and 6 hour after serum stimulation of cilium disassembly. The range of the Log_2_ Fold Change (FC) values is from 0 to 8, represented by shades of purple. Categories were determined using DAVID functional annotation tool and literature mining. Centrosome/cilia/centriolar satellite proteins were plotted in (D) where proteins with both centrosome and centriolar satellite compartment association are shown in bold. Actin-binding proteins were plotted in (E) where proteins shown in bold are ciliary actin-related proteins. Proteins linked to vesicular trafficking and lipid metabolism were plotted in (F) and (G), respectively.

## Notes

### Competing Interest Statement

The authors have declared no competing interest.

